# Reconciling Dimensional and Categorical Models of Autism Heterogeneity: a Brain Connectomics & Behavioral Study

**DOI:** 10.1101/692772

**Authors:** Siyi Tang, Nanbo Sun, Dorothea L. Floris, Xiuming Zhang, Adriana Di Martino, B.T. Thomas Yeo

**Author notes:** Equal contribution (Siyi Tang, Nanbo Sun). **Address correspondence to:** B.T. Thomas Yeo, ECE, CIRC, N.1 & MNP, National University of Singapore.

## Abstract

**Background:** Heterogeneity in autism spectrum disorder (ASD) has hindered the development of biomarkers, thus motivating subtyping efforts. Most subtyping studies divide ASD individuals into non-overlapping (categorical) subgroups. However, continuous inter-individual variation in ASD suggests the need for a dimensional approach.

**Methods:** A Bayesian model was employed to decompose resting-state functional connectivity (RSFC) of ASD individuals into multiple abnormal RSFC patterns, i.e., categorical subtypes henceforth referred to as “factors”. Importantly, the model allowed each individual to express one or more factors to varying degrees (dimensional subtyping). The model was applied to 306 ASD individuals (age 5.2-57 years) from two multisite repositories. Posthoc analyses associated factors with symptoms and demographics.

**Results:** Analyses yielded three factors with dissociable whole-brain hypo/hyper RSFC patterns. Most participants expressed multiple (categorical) factors, suggestive of a mosaic of subtypes within individuals. All factors shared abnormal RSFC involving the default network, but the directionality (hypo/hyper RSFC) differed across factors. Factor 1 was associated with core ASD symptoms, while factor 2 was associated with comorbid symptoms. Older males preferentially expressed factor 3. Factors were robust across multiple control analyses and not associated with IQ, nor head motion.

**Conclusions:** There exist at least three ASD factors with dissociable patterns of whole-brain RSFC, behaviors and demographics. Heterogeneous default network hypo/hyper RSFC across the factors might explain previously reported inconsistencies. The factors differentiated between core ASD and comorbid symptoms - a less appreciated domain of heterogeneity in ASD. These factors are co-expressed in ASD individuals with different degrees, thus reconciling categorical and dimensional perspectives of ASD heterogeneity.

## Introduction

A major challenge in developing biomarkers for Autism Spectrum Disorder (ASD) is the high heterogeneity among ASD individuals. This encompasses core ASD symptoms (1), cognitive skills (2), comorbid psychiatric/medical conditions (3), brain atypicalities (4,5), and genetics (6). Consequently, there have been significant efforts in defining ASD subtypes. Most studies have focused on variability of behavioral or cognitive characteristics (7–10). Studies focusing on brain features are emerging (11–14). Here, we propose a Bayesian framework to decompose whole-brain resting-state functional connectivity (RSFC) patterns in ASD individuals into multiple hypo/hyper RSFC patterns, which we will refer to as “factors” (Figure 1A). This approach allows an individual to express one or more factors (categorical subtypes) to varying degrees (continuous), thus potentially reconciling dimensional (13–15) and categorical (11,12,17) models of ASD heterogeneity.

**Figure 1.**
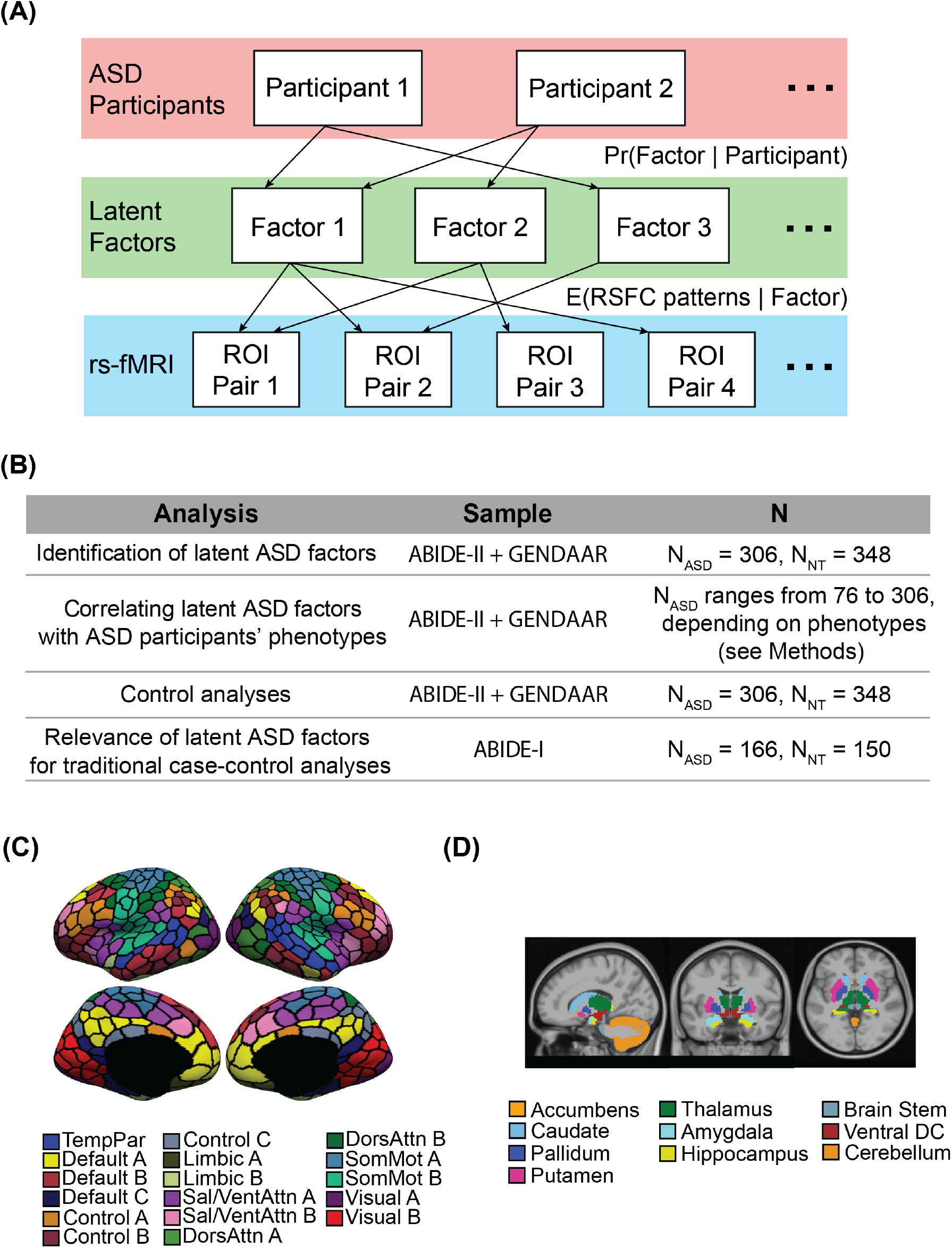
Bayesian model and overview of analyses in this study. **(A)** A Bayesian model of ASD participants, latent factors, and resting-state fMRI (rs-fMRI). The model assumes that each ASD participant expresses one or more latent factors, and each factor is associated with distinct, but possibly overlapping patterns of hypo/hyper functional connectivity. Given the RSFC data of ASD participants and a predefined number of factors K, the model estimates the probability that a participant expresses a latent factor (i.e., factor composition of the participant or Pr(Factor | Participant)), as well as the expected patterns of hypo/hyper RSFC associated with each factor (i.e., factor-specific hypo/hyper RSFC patterns or E(RSFC patterns | Factor)). **(B)** We first identified latent ASD factors using the ABIDE-II+GENDAAR sample. The latent factors were then correlated with ASD participants’ phenotypes (i.e., demographics and behavioral symptoms) in the same sample. Because the multiple sites do not have a uniform set of phenotypes, the sample size varied depending on the phenotype. In addition, we performed several control analyses using the ABIDE-II+GENDAAR sample to ensure the robustness of the latent factors. Finally, we explored the relevance of the latent factors on traditional case-control analyses in the ABIDE-I sample. **(C)** 400 cortical parcels (33). Colors are assigned based on 17 networks widely used in the rs-fMRI literature (34). The 17 networks are divided into eight groups (TempPar, Default, Control, Limbic, Salience/Ventral Attention, Dorsal Attention, Somatomotor and Visual). **(D)** 19 subcortical ROIs (35).

This approach is motivated by two important considerations. First, most previous ASD subtyping studies assumed that each participant belonged to a single (categorical) subtype. By contrast, the term “spectrum” in ASD suggests continuous variation across individuals (18). This is observed at varying degrees across multiple symptom domains (19,20). In parallel, evidence from genetics and neurobiology suggests that autism results from the combination of multiple factors underlying distinct pathways (21,22). Thus, ASD inter-individual variability may reflect different degrees of expression of such factors and related mechanisms (6,23). Together, these observations motivate a mosaic approach to ASD subtyping that incorporates categorical and dimensional features (Figure 1A). Our model allows each ASD individual to express more than one latent factor. For example, the hypo/hyper RSFC pattern of an ASD individual might be explained by 90% factor 1 and 10% factor 2, while the hypo/hyper RSFC pattern of another ASD individual might be explained by 40% factor 1 and 60% factor 2.

Second, early resting-state fMRI (rs-fMRI) investigations supported models of ASD as a dysconnection syndrome (23–26). Although these early studies focused on *a priori* regions/networks of interest in small to moderately sized samples, these observations have been extended by more recent whole-brain investigations of larger samples showing that multiple functional networks subserving the wide range of processes impaired in ASD are affected (26). Importantly, recent studies have reconciled previously inconsistent findings of either hypo- or hyper-connectivity in ASD by showing that both patterns co-exist, though affecting distinct functional circuits (27–29). Nevertheless, these studies rely on traditional case-control analyses, which may miss out on less frequently expressed RSFC patterns due to ASD heterogeneity and/or sampling biases. Thus, our study seeks to provide detailed characterization of the nature and spatial extent of functional dysconnections in ASD, while accounting for significant heterogeneity among ASD individuals.

To address these challenges and estimate latent ASD factors with distinct patterns of whole brain hypo/hyper connectivity, we combined two multisite rs-fMRI data repositories (Autism Brain Imaging Data Exchange-second release; ABIDE-II (31) and the Gender Explorations of Neurogenetics and Development to Advance Autism Research; GENDAAR) (32)). Posthoc analyses were performed to examine common and distinct abnormal RSFC across factors. Furthermore, associations between the latent factors and multiple phenotypic information were examined using multivariate analyses to capture the complexity of ASD.

## Methods and Materials

### Overview

Our analyses proceeded in four steps (Figure 1B). First, to identify latent ASD factors, we applied a Bayesian model (Figure 1A) to a combined dataset comprising ABIDE-II (31) and GENDAAR (32). We used this combined dataset to maximize sample size with regards to both MRI and non-brain-imaging phenotypic data. Second, we examined the associations between latent factors and ASD participants’ phenotypes (i.e., demographic and behavioral symptoms) in the ABIDE-II+GENDAAR combined sample. Third, multiple control analyses were performed to ensure robustness of the results. Lastly, we utilized another independent dataset (ABIDE-I (28)) to explore the drawbacks of case-control analyses, which do not account for ASD heterogeneity.

### Participants

MRI data selected from the ABIDE and GENDAAR repositories were analyzed. Following preprocessing and quality control (see “MRI preprocessing” and Supplemental Methods), the resulting sample comprised 242 ASD and 276 neurotypical (NT) participants from ABIDE-II, which were combined with 64 ASD and 72 NT participants from GENDAAR for primary analyses, as well as an independent sample of 166 ASD and 150 NT participants from ABIDE-I (secondary analyses). Age, sex and head motion were matched between ASD and NT participants within each site. Participants’ characteristics are summarized in Tables 1 and S1.

**Table 1.**
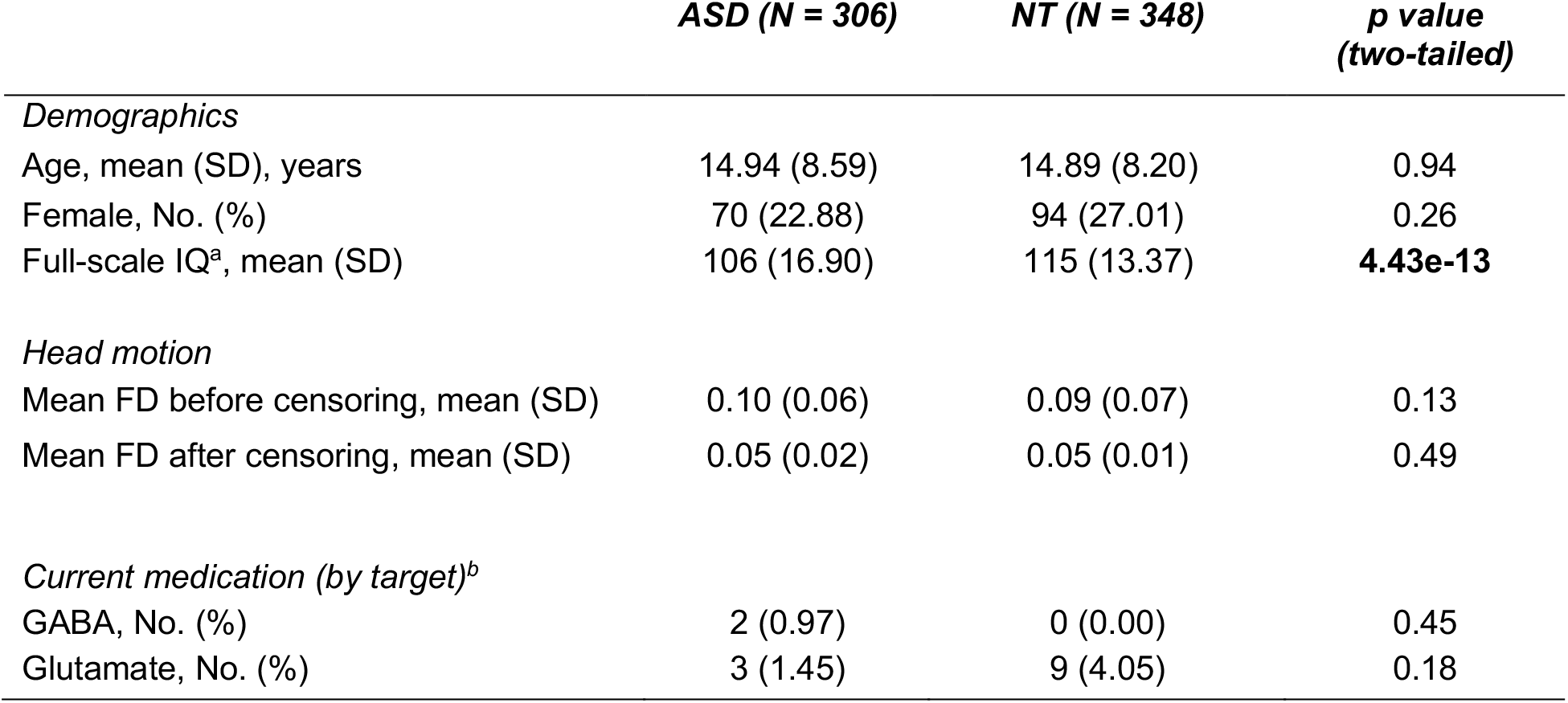
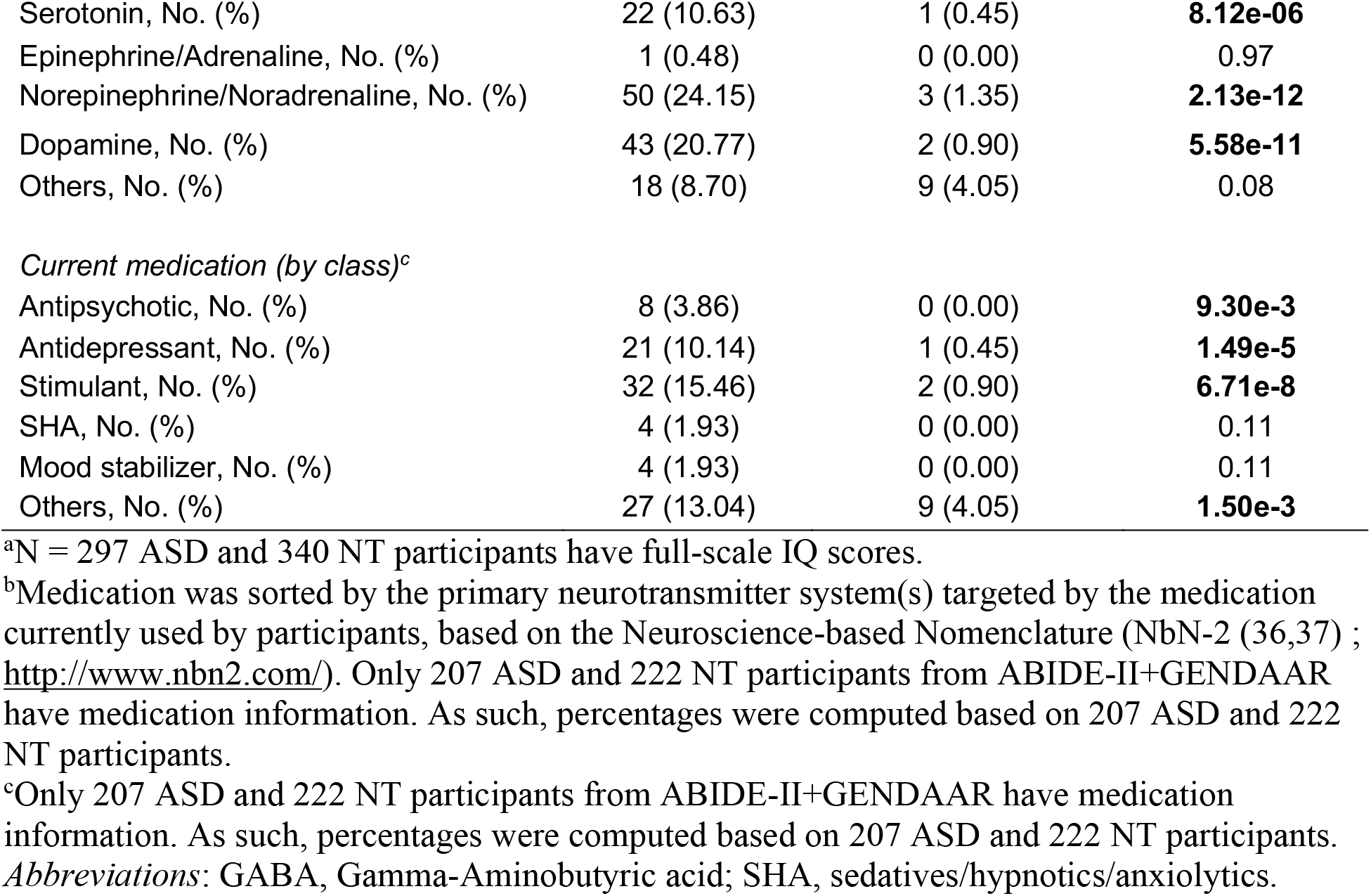
Characteristics and behavioral data of participants from ABIDE-II and GENDAAR. Fourteen data collections from ABIDE-II and GENDAAR were included in this study: BNI_1 (N_ASD_ = 21, N_NT_ = 15), ETH_1 (N_ASD_ = 6, N_NT_ = 21), GU_1 (N_ASD_ = 34, N_NT_ = 26), IP_1 (N_ASD_ = 10, N_NT_ = 5), IU_1 (N_ASD_ = 17, N_NT_ = 15), KKI_1 (N_ASD_ = 31, N_NT_ = 90), NYU_1 (N_ASD_ = 42, N_NT_ = 23), OHSU_1 (N_ASD_ = 23, N_NT_ = 23), TCD_1 (N_ASD_ = 14, N_NT_ = 18), UCD_1 (N_ASD_ = 15, N_NT_ = 9), UCLA_1 (N_ASD_ = 11, N_NT_ = 11), USM_1 (N_ASD_ = 10, N_NT_ = 13), U_MIA_1 (N_ASD_ = 8, N_NT_ = 7), GENDAAR (N_ASD_ = 64, N_NT_ = 72). ASD and NT participants were compared with either two-sample t-tests (for continuous measures) or chi-squared tests (for categorical measures). All p-values that survived false discovery rate (FDR) correction (q < 0.05) are indicated in bold.

### MRI preprocessing

The neuroimaging data were processed using a previously published pipeline (38–40). See Supplemental Methods for details. Here, we briefly outline the procedure. Rs-fMRI data underwent standard preprocessing including slice time correction, motion correction and alignment with anatomical T1. Frame-wise displacement (FD) and voxel-wise differentiated signal variance (DVARS) were computed using fsl_motion_outliers (41,42). Volumes with FD > 0.2 mm or DVARS > 50 were marked as censored frames, together with one frame before and two frames after. Uncensored data segments lasting fewer than five contiguous volumes were also censored (43). Functional runs with more than 50% censored frames were removed (Table S2).

We regressed out 18 nuisance regressors, consisting of six motion parameters, averaged cerebrospinal ventricular signal, averaged white matter signal, global signal and their temporal derivatives. Censored frames were ignored when regression coefficients were computed. Data were interpolated across censored frames using least squares spectral estimation (44). We chose to regress the global signal because of its effectiveness in removing motion-related and respiratory artifacts (45,46). Recent work has also suggested that global signal regression (GSR) increases the associations between behavior and RSFC (39). Nevertheless, we performed control analyses using an alternative to GSR (see “Control analyses”). Finally, the data were band-pass filtered (0.009Hz ≤ f ≤ 0.08Hz), projected onto FreeSurfer fsaverage6 surface space, smoothed with a 6mm kernel and down-sampled onto FreeSurfer fsaverage5.

### Resting-state functional connectivity (RSFC)

We utilized a cortical parcellation (33) comprising 400 cortical regions-of-interest (ROIs; Figure 1C) and a subcortical segmentation (35) comprising 19 subcortical ROIs (Figure 1D). RSFC (Pearson’s correlation) was computed among the average time series of 419 brain ROIs (ignoring censored frames), yielding a 419 × 419 RSFC matrix for each participant. Age, sex, head motion (mean frame-wise displacement (42)) and site differences were regressed out from all participants’ RSFC data with a general linear model (GLM). Regression coefficients were estimated only from NT participants to retain any ASD-specific interactions with participants’ characteristics (e.g., age). Each RSFC entry (i.e., lower triangular entries since the matrices are symmetric) of the ASD participants was z-normalized with respect to the 348 ABIDE-II+GENDAAR NT participants. A z-score larger (or smaller) than zero for a given ROI pair indicates hyper-connectivity (or hypo-connectivity) relative to the NT participants.

### Latent factors in ABIDE-II+GENDAAR

Latent ASD factors were identified using the ABIDE-II+GENDAAR dataset. We applied a Bayesian model (Figure 1A) to the z-normalized RSFC of the ASD participants to estimate latent factors. The model is a variant of Bayesian models previously utilized to discover latent atrophy factors in Alzheimer’s Disease (47) and latent components subserving cognitive tasks (48). It assumes that each individual expresses one or more latent factors, associated with distinct, but possibly overlapping patterns of hypo/hyper RSFC. Given the RSFC data and a user-defined number of factors *K*, we can estimate the factor composition of each participant, i.e., probability that a participant expresses a latent factor (i.e., Pr(Factor | Participant)), as well as the factor-specific hypo/hyper RSFC patterns (i.e., E(RSFC patterns | Factor)). See Supplemental Methods.

We estimated 2, 3 and 4 latent factors. The estimations were robust for 2 and 3 factors, but unstable for 4 factors (Figure S1). Therefore, a larger number of factors was not considered. Furthermore, the two-factor estimates were inconsistent in the control analyses (see “Control analyses”), so we focused on the three-factor estimates in subsequent analyses.

To estimate confidence intervals for the factor-specific hypo/hyper RSFC patterns, we applied a bootstrapping procedure that generated 100 samples from ASD participants’ z-normalized RSFC data. Z-scores were then calculated by dividing factor-specific hypo/hyper RSFC patterns by the bootstrap-estimated standard deviation. To reduce multiple comparisons, the factor-specific hypo/hyper RSFC patterns were averaged across ROI pairs within and between the 17 networks and subcortical structures (Figures 1C-1D), resulting in 18 × 18 matrices, before computing bootstrapped z-scores. The z-scores were converted to p-values and corrected using false discovery rate (q < 0.05) along with other tests (Supplemental Methods).

### Associations between participants’ characteristics and latent factors in ABIDE-II+GENDAAR

We applied separate GLM (or logistic regression for binary variables) to the factor compositions and each characteristic (age, sex, FIQ and head motion) of ABIDE-II+GENDAAR ASD participants to investigate potential associations. For each GLM/logistic regression, participants’ characteristic and factor compositions were treated as the dependent and independent variables respectively (Supplemental Methods).

### Associations between behavioral symptoms and latent factors in ABIDE-II+GENDAAR

Because ABIDE-II+GENDAAR consisted of datasets across independent sites, not all participants had the same behavioral measures (Table S3). If we considered all available behavioral measures jointly, we would be left with only seven participants. Therefore, available behavioral scores were divided into five groups to maximize the number of participants in each group. For example, Social Responsiveness Scale (SRS) Autistic Mannerism and Repetitive Behaviors Scale-Revised 6 Subscales (RBSR-6) subscales were grouped together because they index aspects of restricted/repetitive behaviors (RRB). Of note, Autism Diagnostic Observation Schedule (ADOS) Stereotyped Behavior subscore was not included into the RRB domain because only 38 participants had ADOS Stereotyped Behavior with SRS Autistic Mannerism, RBSR-6 subscales. The five groups of behavioral scores are shown in Table S4.

We then applied canonical correlation analysis (49) (CCA) between each group of behavioral scores and each factor loading (i.e., Pr(Factor | Participant)), i.e., fifteen CCAs in total in the case of the three-factor model (see Supplemental Methods). The goal of the CCA was to find an optimal linear combination of the behavioral scores that maximally correlated with the factor loading. Age, sex, head motion and sites were regressed out from both behavioral scores and factor loadings before the CCA. Statistical significance was tested using 10,000 permutations that accounted for different sites. False discovery rate (q < 0.05) was utilized to correct for multiple comparisons (Supplemental Methods).

### Control analyses in ABIDE-II+GENDAAR

We performed several control analyses to ensure robustness of results. First, to ensure robustness to preprocessing strategies, we applied the Bayesian model to rs-fMRI processed using CompCor (50) instead of GSR. Second, we applied k-means clustering to ABIDE-II+GENDAAR ASD participants’ z-normalized RSFC data (processed with GSR or CompCor) to ensure robustness to analysis strategies (k-means versus Bayesian model). Third, we compared behavioral associations of the k-means clusters with those of the latent factors. Lastly, we randomly split the 306 ASD participants in ABIDE-II+GENDAAR into two groups (Table S5) and estimated the latent factors in each group independently. See Supplemental Methods for details.

### Drawbacks of traditional case-control analyses in ABIDE-I

In most studies, ASD and NT individuals are compared without accounting for ASD heterogeneity. To explore the drawbacks of case-control analyses, we considered 166 ASD and 150 NT participants from ABIDE-I (Table S1) and inferred their factor compositions using latent factors estimated from ABIDE-II+GENDAAR. These compositions were used to assign each individual to one of three subgroups. This sub-grouping violates the spirit of our hybrid dimensional-categorical approach, but is necessary for comparison with traditional case-control analysis. To ensure robustness, we experimented with two different criteria of assigning ASD participants to subgroups (Tables S6-S7). RSFC differences between each ASD subgroup and demographically-matched NT participants were computed and compared with traditional case-control analysis (i.e., RSFC differences between ASD and NT participants without subgrouping). See Supplemental Methods for details.

## Results

### Latent ASD factors with dissociable hypo/hyper RSFC patterns

We applied the Bayesian model (Figure 1A) to 306 ABIDE-II+GENDAAR ASD participants. An important model parameter is the number of latent factors *K*. We experimented with *K* = 2, 3 and 4. Four-factor model was unstable (Figure S1), so we did not explore more factors. On the other hand, the two-factor model was sensitive to the preprocessing strategy (Supplemental Results; Table S8). Thus, we focused on the three-factor solution.

Each of the three factor-specific hypo/hyper RSFC patterns among the 400 cortical and 19 subcortical ROIs (Figures 1C-1D) are shown in Figure 2A (unthresholded) and Figure 2B (statistically significant). Figure 2C illustrates the significant RSFC patterns averaged within and between the 17 networks and subcortical structures.

**Figure 2.**
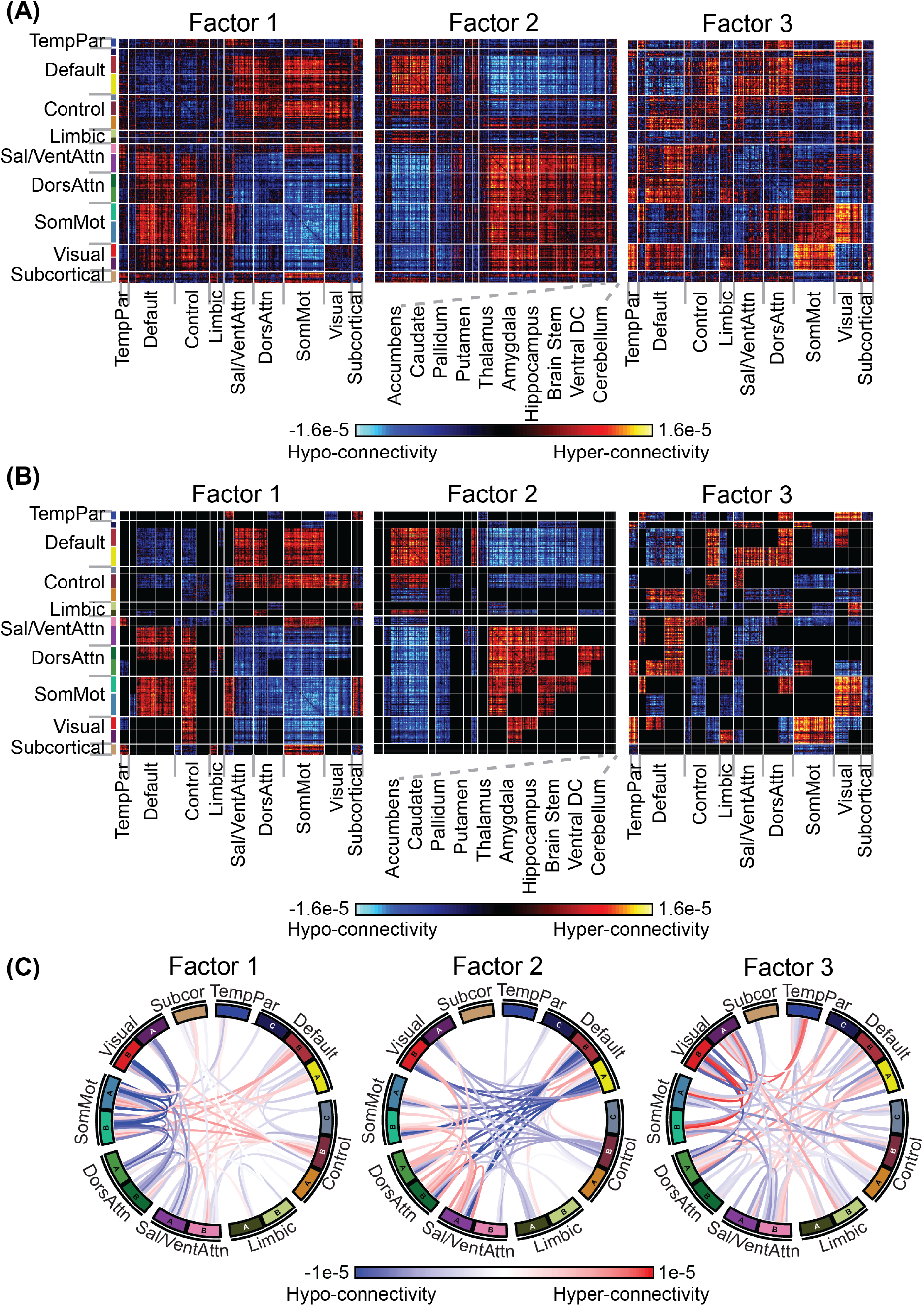
Three latent ASD factors estimated from ABIDE-II+GENDAAR ASD participants. **(A)** Patterns of hypo/hyper RSFC (unthresholded) associated with each factor. Hot color indicates hyper-connectivity (relative to NT), and cold color indicates hypo-connectivity (relative to NT). **(B)** Statistically significant patterns of hypo/hyper RSFC associated with each factor. **(C)** Significant patterns of hypo/hyper RSFC associated with each factor, averaged within and between networks.

Factor 1 was associated with ASD-related hypo-connectivity (blue in Figure 2) within and between perceptual/motor networks (somatomotor A/B, visual A/B, salience/ventral attention A, dorsal attention A/B). On the other hand, there was ASD-related hyper-connectivity (red in Figure 2) between perceptual/motor and association networks (default, control and salience/ventral attention B), as well as between somatomotor and subcortical regions (caudate and thalamus).

Factor 2 was associated with a pattern of hypo/hyper RSFC almost opposite to factor 1 (r = −0.57), but with subtle deviations. For example, regions within default networks A and B were strongly hyper-connected in factor 2, but only weakly hypo-connected in factor 1. Similarly, regions between somatomotor networks and caudate were strongly hyper-connected in factor 1, but did not exhibit any atypical RSFC in factor 2.

Factor 3 was characterized by a complex pattern of hypo/hyper RSFC. For example, there was hyper-connectivity between visual and somatomotor networks. There was also strong hypo-connectivity among regions within default networks A/B, and among regions within the visual networks.

### Factor compositions of ASD participants in ABIDE-II+GENDAAR

Figure 3 shows the factor compositions of ASD participants in ABIDE-II+GENDAAR. Most participants expressed multiple latent factors rather than a single factor. Critically, no single site showed predominantly one single factor, suggesting that latent factors were not driven by site differences.

**Figure 3.**
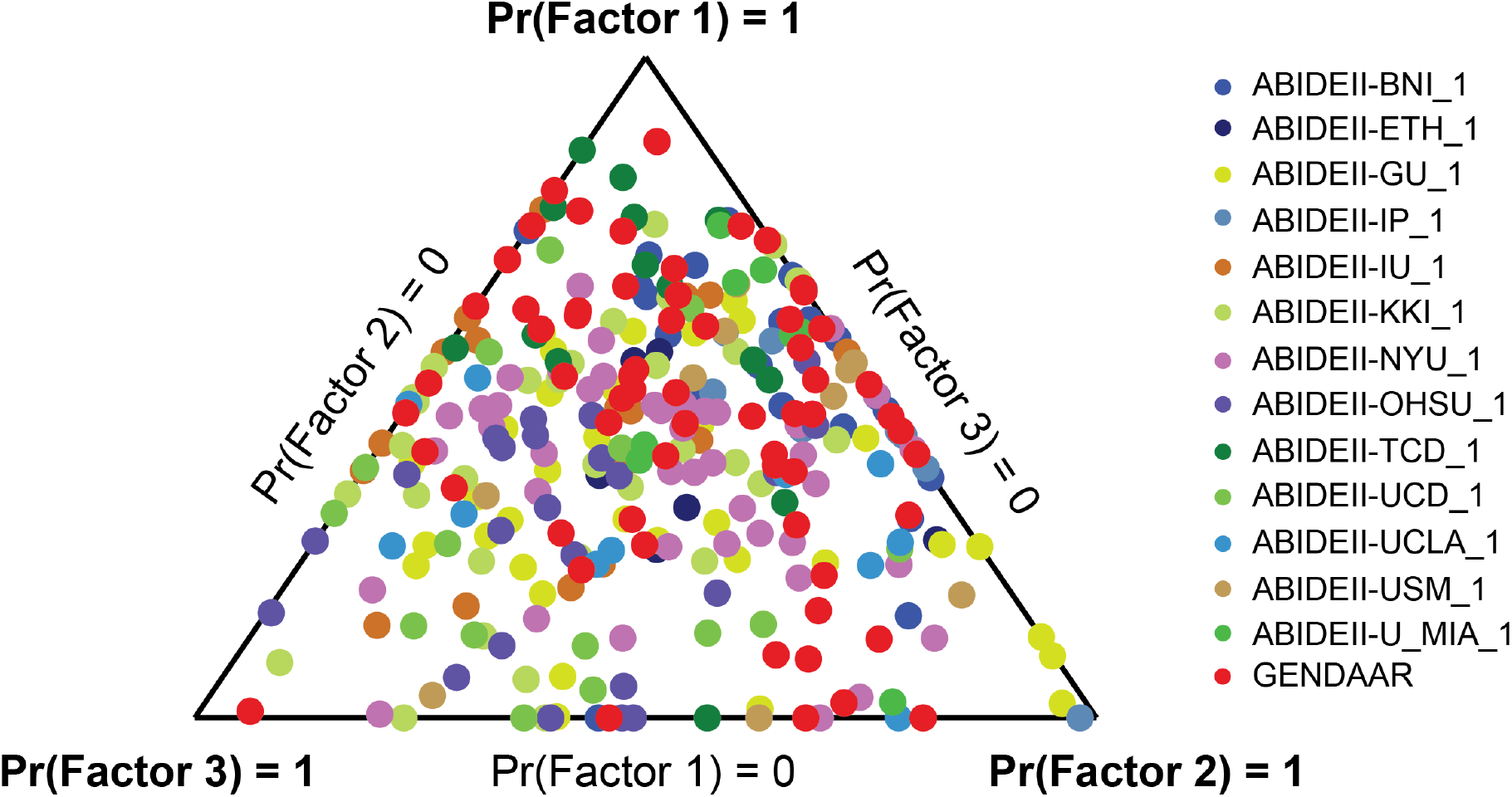
Factor compositions of ASD participants in ABIDE-II+GENDAAR. Each participant corresponds to a dot, with location (in barycentric coordinates) representing the factor composition (i.e., Pr(Factor | Participant)). Corners of the triangle represent pure factors, and dots closer to the corner indicates higher probability of the respective factor. Most dots are far from the corners, suggesting that most ASD participants expressed multiple factors.

### Default network exhibits abnormal connectivity across all three factors

By comparing the RSFC patterns among the three factors (Figure 2B), Figure S2 shows the statistically significant hypo/hyper RSFC unique to each latent factor.

On the other hand, to examine hypo/hyper RSFC patterns that are shared across factors, within-network and between-network blocks with significant bootstrapped z-scores (Figure 2C) were binarized (ignoring directionality of abnormality) and summed across the three factors (Figure 4A). In addition, absolute values of hypo/hyper RSFC patterns that were significant across all three factors (Figure 2B) were summed to obtain the magnitude of hypo/hyper RSFC patterns common across factors (Figure 4B). Altered connectivity within default A and B networks were notable, as well as between default and perceptual/motor networks (somatomotor A, salience/ventral attention A, dorsal attention B). In addition, hypo/hyper connectivity within salience/ventral attention A, within dorsal attention, as well as between somatomotor and control B networks were also common across all three factors.

**Figure 4.**
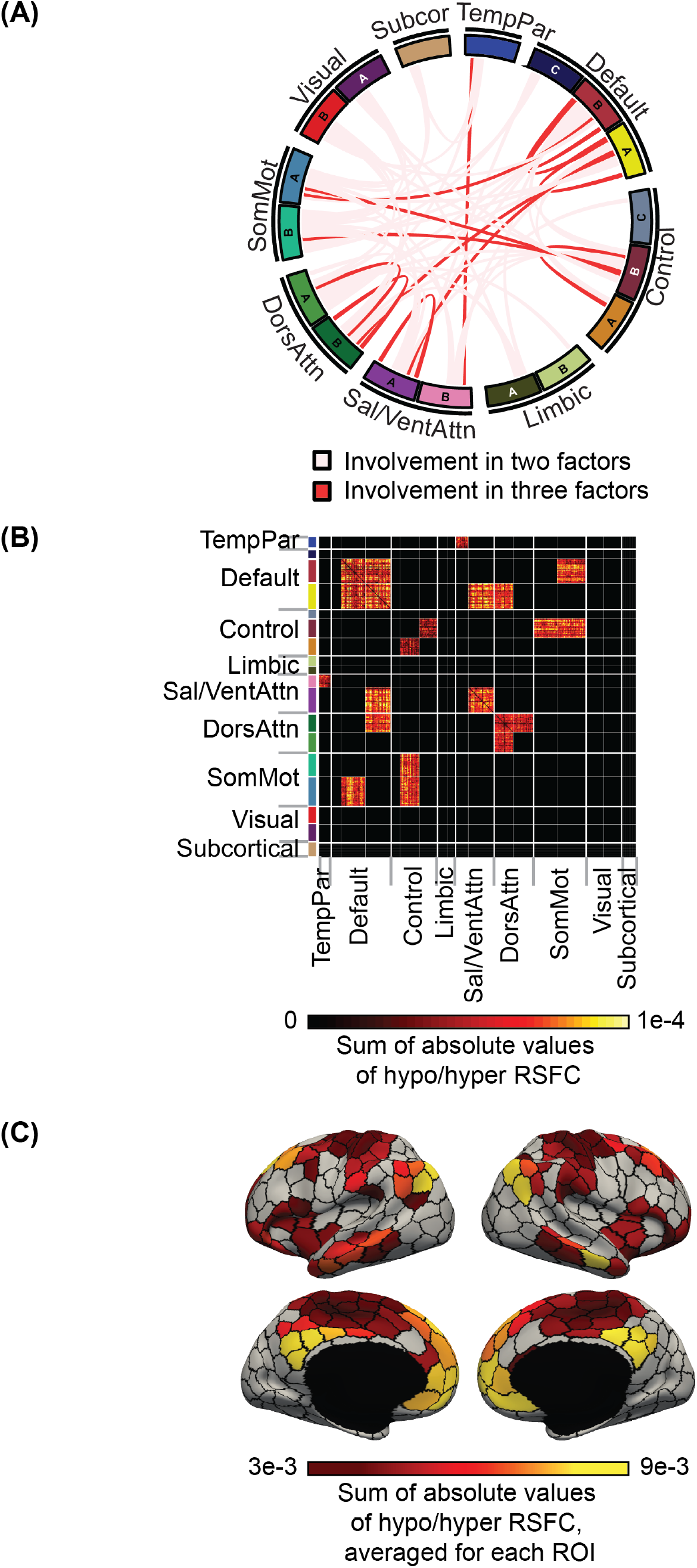
Patterns of hypo/hyper RSFC involved in all three factors. **(A)** Statistically significant within- and between-network hypo/hyper RSFC patterns (Figure 2C) were binarized, and then summed across the three factors; hypo/hyper RSFC patterns that were significant in only one factor were set to zero. **(B)** Sum of absolute values of hypo/hyper RSFC (Figure 2B) that were significant across all three factors. **(C)** The summed absolute values of hypo/hyper RSFC from panel (B) were averaged across the rows for each ROI, and projected onto a surface map for visualization.

Lastly, Figure 4C shows the strength of involvement of each ROI obtained by summing the rows of Figure 4B. The strength of default network’s involvement was particularly striking. Although atypical default network connectivity was present across all factors, the directionalities were inconsistent. For example, factor 2 exhibited hyper-connectivity within the default network, while factor 1 and 3 exhibited hypo-connectivity (Figure 2B).

### Participants’ characteristics across latent factors in ABIDE-II+GENDAAR

We used GLM (or logistic regression) to investigate whether ASD participants’ characteristics (i.e., age, sex, FIQ, head motion) varied across factors in ABIDE-II+GENDAAR. Factor 3 was more strongly expressed by males relative to factors 2 (*p*<0.001; Figure S3A). Factor 3 was also associated with older participants compared to factors 1 and 2 (*p*=0.002 and *p*=0.01 respectively; Figure S3B). There was no difference in FIQ nor head motion across factors (Figure S3C and S3D).

### Associations between latent factors and behavioral symptoms

To examine associations between latent factors and behavioral symptoms in ABIDE-II+GENDAAR ASD participants, we performed CCA between each factor loading and each group of behavioral scores (see Methods). Four sets of CCA analyses remained significant after FDR correction (q < 0.05; Figure 5). Higher scores indicated worse symptoms. Therefore, positive values implied that a higher loading on the factor was associated with greater impairment. Factor 1 was associated with both worse RRB (Figure 5A; *r*=0.54, *p*=0.002) and social deficits (Figure 5B; *r*=0.27, *p*=0.004). Factor 2 was associated with both worse externalizing problems (Figure 5C; *r*=0.43, *p*=0.004) and executive dysfunction (Figure 5D; *r*=0.33, *p*=0.02). Factor 3 was not associated with any aggregated behavioral symptoms.

**Figure 5.**
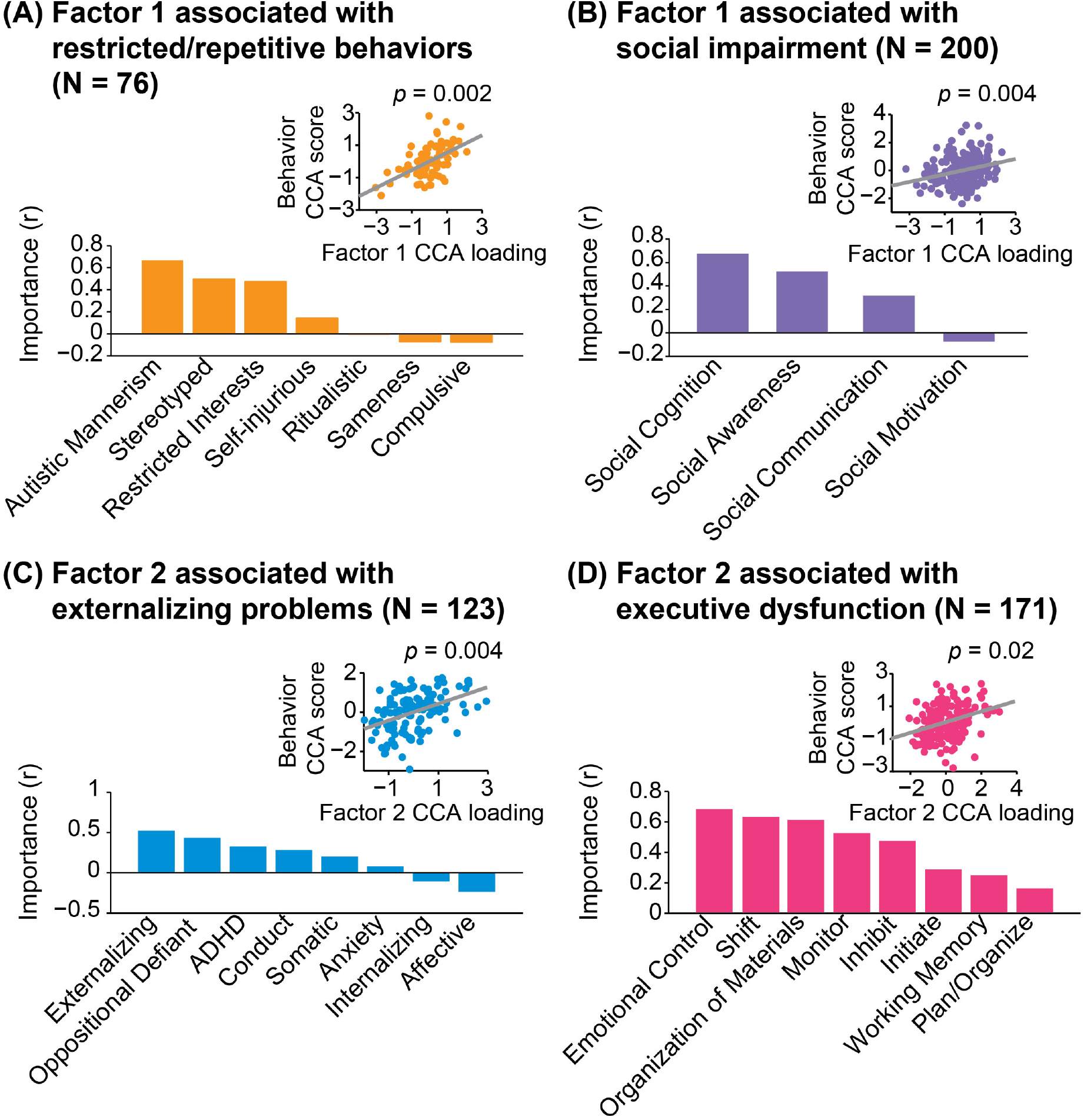
First and second latent ASD factors were associated with distinct behavioral deficits. CCA analyses between the loadings of each factor and five groups of behavioral scores revealed that RSFC factors 1 and 2 were associated with distinct behavioral deficits. Four sets of CCA analyses remained significant after FDR (*q* < 0.05) multiple comparisons correction. **(A)** Associations between factor 1 and RRB measured by SRS Autistic Mannerism subscale and RBSR-6 subscales. **(B)** Associations between factor 1 and social responsiveness measured by SRS subscales (excluding SRS Autistic Mannerism). **(C)** Associations between factor 2 and comorbid psychopathology measured by CBCL-6-18 subscales. **(D)** Associations between factor 2 and executive function measured by BRIEF subscales. The bar plots show the Pearson’s correlation between each behavioral score and CCA behavioral loading (more details in Supplemental Methods). Positive correlation suggests that a higher loading on the factor was associated with greater impairment. The scatterplots show the relationship between CCA behavioral score loading and CCA ASD factor loading, where each dot represents an ASD participant. Factor 1 was associated with worse RRB and social deficits, while factor 2 was associated with worse externalizing problems and executive dysfunction. *Abbreviations*: SRS, Social Responsiveness Scale; RBSR-6, Repetitive Behaviors Scale-Revised 6 Subscales; CBCL-6-18, Child Behavior Checklist Ages 6-18; BRIEF, Behavior Rating Inventory of Executive Function. See Table S4 for behavioral scores entered into the CCA.

### Control analyses

Here, we summarize the results of the control analyses (see Supplemental Results for more details). First, factors 1 and 2 from the three-factor model were similar regardless of processing pipeline (GSR or CompCor), but not so for factor 3 (Table S8). On the other hand, all three factors (estimated in primary analyses using GSR; Figure 2) were similar to clusters obtained from k-means regardless of GSR or CompCor (Table S9). Thus, overall, factors 1 and 2, and to some extent factor 3, were relatively robust to preprocessing and analysis strategies.

Compared to the latent factors, k-means clusters showed similar but weaker behavioral associations (Figure S4), suggesting potential advantage of our hybrid dimensional-categorical model. Finally, factors estimated from random splits of ABIDE-II+GENDAAR ASD participants were similar to the factors generated in primary analyses and with each other (Table S10).

### Traditional case-control analysis yields smaller effects and misses significant ASD-related RSFC associations

To explore potential drawbacks of traditional case-control analyses, we computed RSFC differences between 166 ASD and 150 demographically-matched NT participants from ABIDE-I. We also computed RSFC differences between ASD and NT participants from each ASD subgroup in ABIDE-I (see Methods). Despite the larger sample size, traditional case-control analysis yielded significantly weaker RSFC differences than the subgroup analyses and missed out on ASD-related RSFC differences (Figures S5-S6). See Supplemental Results for details.

## Discussion

In this study, we applied a Bayesian model to a large rs-fMRI cohort of individuals with ASD, revealing three latent factors with dissociable patterns of hypo/hyper RSFC. Each factor was expressed to different degrees across individuals and associated with distinct behavioral and demographic features that are known sources of clinical heterogeneity in ASD, i.e., core ASD impairments, externalizing symptoms, executive dysfunctions, age and sex. Overall, these results suggest that each ASD individual expresses a mosaic of latent factors. This has been missed in prior subtyping approaches that assigned each individual to a single subtype (7–12,51). By contrast, our approach allows each individual’s factor composition to be unique, thus retaining inter-individual variability. This is consistent with models suggesting that ASD heterogeneity reflects the contribution of multiple mechanisms with different degrees of expression across subjects (6,23).

Factor 1 loading was the highest when averaged across ASD participants, so unsurprisingly, the hypo/hyper RSFC pattern of factor 1 was the most similar to prior case-control whole-brain comparisons emerging from distinct analytical approaches applied to partially overlapping samples from ABIDE-I (27–29). ASD-related hypo-connectivity within sensory and salience networks, as well as hyper-connectivity between somatomotor and subcortical regions were particularly notable. Likely reflecting its greater prevalence among ASD participants, this factor was more strongly associated with the core ASD symptoms of both social reciprocity and RRB (20). Reflecting their wide range of ASD symptoms, factor 1 involved multiple functional networks previously associated with ASD severity. For example, social skills impairment and hypo-RSFC within default or salience networks have been previously reported (52,53). Although not often explored, hyper-connectivity between thalamus and temporal cortex have also been associated with social reciprocity deficits in ASD (30), and an atypical RSFC balance in cortical striatal circuitry involving limbic somatomotor and frontoparietal networks have been associated with RRB indexed by RBS-R total scores (54).

Interestingly, the factors did not differentiate among core ASD symptoms (e.g., RRB versus social reciprocity), but differentiated between core ASD symptoms (factor 1) and comorbid symptoms (executive dysfunction and externalizing symptoms; factor 2). This finding underscore two important aspects of the ASD phenomenology. One is the strong intercorrelation of ASD symptom domains (20,55) and possibly their partially overlapping biological underpinnings. While neuroimaging studies with deeper phenotyping may be able to characterize the association between the RSFC patterns in a given factor and specific symptom subdomains, larger scale phenomena reflected in the mosaic of atypical RSFC in a given factor are at play. This is suggested by recent reports of atypical hierarchical network organization in ASD relative to controls, which was also related to global metrics of ASD traits (56,57). The second aspect of the ASD phenomenology underscored by our brain-behavioral findings is that comorbidity critically contributes to ASD heterogeneity. This is a clinical dimension that has been quantitatively and qualitatively overlooked in the neuroimaging literature, albeit with notable exceptions (e.g., 56,57). Our findings suggest that comorbidity should be more consistently accounted for in ASD biomarker efforts.

In contrast with factor 1, factor 2 was characterized by a pattern of hypo/hyper RSFC that included, but was not limited to, hyper-connectivity within the default and attentional networks, as well as hypo-connectivity between the default and attentional networks. As previously mentioned, factor 2 was also associated with executive dysfunction and externalizing symptoms. These findings are consistent with reports that poor executive control might result from abnormalities in attentional networks (60). Greater connectivity within default network has also been linked to worse executive function in ASD (53). Furthermore, externalizing symptoms are frequently observed in large proportions of ASD individuals (59–61). Finally, it remains unknown to what extent factor 2, as well as the other factors and their relationship with behavior are ASD specific or extend across diagnoses. An initial study (13) has suggested that shared factors across ADHD and ASD might exist, but the lack of a shared behavioral battery between the ADHD and ASD samples limited further exploration with specific symptom domains. Another study has shown that distinct clusters of RSFC patterns exist across individuals with and without ASD, and the subtypes revealed unique brain-behavior relations (14). The emergence and availability of deeply-phenotyped transdiagnostic samples (64) will help address this gap.

In our study, factors 1 and 2 were associated with similar age, but distinct behavioral deficits, suggesting that they might not simply reflect disease severity or neurodevelopment stage. On the other hand, factor 3 was more frequently expressed in older participants, and might thus reflect a neurodevelopmental stage. The prevalence of ASD is much higher in males than females (65). Recent studies have suggested that ASD-related sex differences are associated with atypical brain connectivity and mentalization (65–67). Our study suggests that factor 3 was associated with male participants, but given the small percentage of female participants (23%) in ABIDE-II+GENDAAR, replication with larger sex-balanced datasets is necessary.

Finally, along with factor-specific RSFC signatures, all three factors shared atypical RSFC involving the default network. This is consistent with the larger RSFC ASD literature that have consistently reported structural, and functional ASD-related abnormalities involving the default network (68–70). The directionality of the abnormal default network RSFC varied by factor, e.g., factor 2 exhibited hyper-connectivity within the default network, while factor 1 and 3 exhibited hypo-connectivity (Figure 2B). Similarly, factor 2 exhibited hypo-connectivity between the default and attentional networks, but factors 1 and 3 exhibited hyper-connectivity. The differences between factors may explain some of the inconsistent reports in prior studies on the nature of the DN atypicalities in ASD (50,71–73). As shown by our results from additional analyses on the independent ABIDE-I sample, common case-control comparisons fail to appreciate heterogeneous ASD-related RSFC abnormalities, overall urging for future quantitative ASD subtyping efforts.

A major limitation of this study is the lack of uniform deep phenotyping across all ASD participants because the datasets were gathered post-hoc. Therefore, despite the large sample size, the behavioral associations were separated into five sets of analyses performed on overlapping subsets of ASD participants. A harmonized dataset would also allow the identification of latent factors from RSFC and behavioral deficits simultaneously using a multi-modal variant of the current approach (75). Second, since medication information is limited (e.g., only 8 ASD participants in ABIDEII+GENDAAR were reported to be taking antipsychotic medications), we did not explore the association between factors and medications. Finally, longitudinal data is needed to clarify whether the latent factors reflect ASD heterogeneity or neurodevelopmental stages or complex interactions between heterogeneity and neurodevelopment.

## Conclusion

Our study revealed three latent ASD factors with dissociable whole-brain hypo/hyper RSFC patterns. The factors were associated with distinct behavioral symptoms (core ASD versus comorbid symptoms) and demographics. Our approach allows each individual to express multiple latent factors to varying degrees, rather than a single factor. Therefore, each individual’s factor composition is unique, which might be potentially useful for future biomarker development.

## Disclosures and Acknowledgments

The authors report no competing interests. This manuscript has been posted on bioRxiv.

This work was supported by Singapore MOE Tier 2 (MOE2014-T2-2-016), NUS Strategic Research (DPRT/944/09/14), NUS SOM Aspiration Fund (R185000271720), Singapore NMRC (CBRG/0088/2015), NUS YIA and the Singapore National Research Foundation (NRF) Fellowship (Class of 2017) and partially supported by NIMH R01MH105506-01. Our research also utilized resources provided by the Center for Functional Neuroimaging Technologies, P41EB015896 and instruments supported by 1S10RR023401, 1S10RR019307, and 1S10RR023043 from the Athinoula A. Martinos Center for Biomedical Imaging at the Massachusetts General Hospital. Our computational work was partially performed on resources of the National Supercomputing Centre, Singapore (https://www.nscc.sg). We thank the investigators and sites who contributed to the ABIDE repositories. Details on the funding sources of each site can be found at http://fcon_1000.projects.nitrc.org/indi/abide/. We also thank the GENDAAR Consortium for their efforts in data collection and sharing. The consortium includes, in alphabetical order, Elizabeth H. Aylward, Raphael A. Bernier, Susan Y. Bookheimer, Mirella Dapretto, Nadine Gaab, Daniel H. Geschwind, Andrei Irimia, Allison Jack, Charles A. Nelson, Kevin A. Pelphrey, Matthew W. State, John D. Van Horn, Pamela Ventola and Sara J. Webb. Data from GENDAAR Consortium were obtained from the NIH-supported National Database for Autism Research (NIMH Data Repositories Study No. 2021; NIMH grant R01 MH100028; Principal Investigator: Kevin A. Pelphrey). National Database for Autism Research (NDAR; https://ndar.nih.gov/) is a collaborative informatics system created by the National Institutes of Health to provide a national resource to support and accelerate research in autism. This manuscript reflects the views of the authors and may not reflect the opinions or views of the NIH or of the submitters submitting original data to NDAR.

