## Supplemental materials for "Reconciling Dimensional and Categorical Models of Autism Heterogeneity: a Brain Connectomics & Behavioral Study"

#### Supplemental Methods

Sections S1, S2 and S3 provide more details about participant inclusion criteria, MRI preprocessing and functional connectivity analyses, respectively. Sections S4 and S5 provide more details about the Bayesian model and estimation procedure. Sections S6 and S7 elaborate on the procedures to establish associations between latent factors and participants' characteristics (e.g., age) or behavioral symptoms. Sections S8 and S9 discuss the control analyses to examine the robustness of our results. Section S10 details the analyses to explore drawbacks of traditional case-control analyses. Finally, sections S11 and S12 discuss multiple comparisons correction and code/data availability.

##### S1. Participant inclusion criteria

We selected participants from the Autism Brain Imaging Data Exchange-I (1) (ABIDE-I), ABIDE-II (2) and GENDAAR (3) repositories. Participant inclusion criteria were as follows: (a) the participant had both T1-weighted image and resting-state fMRI (rs-fMRI) scan covering most of the cerebellum; (b) the participant had at least one functional run remaining after rs-fMRI pre-processing and additional quality control described in the next section; (c) the participant's full-scale IQ (FIQ) was not an outlier in the respective sample (i.e. ABIDE-II+GENDAAR or ABIDE-I; FIQ values beyond 1.5 inter-quartile range in the sample were considered as outliers; one ABIDE-II NYU_1 participant, four ABIDE-II IP_1 and one ABIDE-I YALE participants were excluded); (d) the acquisition site had at least five ASD and five NT participants meeting the above criteria. A subset of NT participants was then selected in order to group-match the remaining ASD participants by age, sex and head motion (as measured by fsl_motion_outliers) for each site. The final sample consisted of 166 ASD and 150 neurotypical (NT) participants from ABIDE-I, 242 ASD and 276 NT participants from ABIDE-II, and 64 ASD and 72 NT participants from GENDAAR. Participants' characteristics are summarized in Table 1 and Table S1.

##### S2. Processing of MRI data

The neuroimaging data were processed using a previously published pipeline (4-6). The pipeline code is publicly available here: https://github.com/ThomasYeoLab/CBIG/tree/master/stable_projects/preprocessing/CBIG_fMRI_Preproc2016.

Structural MRI data were processed using FreeSurfer 5.3.0 (http://surfer.nmr.mgh.harvard.edu), which provides automatic algorithms for cortical reconstruction and volumetric segmentation from individual subject's T1 images (7-9). Each subject's cortical surface mesh was registered to a common spherical coordinate system (10,11).

Rs-fMRI data were pre-processed with the following steps: (a) removal of first 4 frames; (b) slice time correction with FSL package (12,13); (c) motion correction using rigid body translation and rotation with FSL package; (d) alignment with structural image using boundary-based registration (14) in FsFast (http://surfer.nmr.mgh.harvard.edu/fswiki/FsFast). Quality control was done by visual inspection of cortical reconstruction and alignment of structural and functional images. Functional runs with boundary-based registration costs (14) larger than 2 standard deviations above the sample mean were excluded.

Frame-wise displacement (FD) and voxel-wise differentiated signal variance (DVARS) were computed using fsl_motion_outliers (12,13). Volumes with FD > 0.2 mm or DVARS > 50 were marked as outliers (i.e., censored frames). One frame before and two frames after the censored frames were also flagged as censored. Uncensored data segments lasting fewer than five contiguous volumes were also labeled as censored (15). Functional runs with more than half of the volumes labeled as censored were removed. The number of participants excluded in this censoring step is shown in Table S2.

Linear regression of multiple nuisance variables was applied. Nuisance regressors consisted of (a) a vector of ones and linear trend; (b) six motion correction parameters; (c) averaged white matter signal; (d) averaged ventricular signal; (e) global signal; (f) temporal derivatives of (b)-(e). The censored frames were ignored when regression coefficients were computed. The data were interpolated across censored frames using least squares spectral estimation of the values at censored frames (16). We chose to regress global signal because of its effectiveness in removing motion-related and respiratory artifacts (17,18). Furthermore, recent work has shown that global signal regression (GSR) greatly increased the associations between behavior and RSFC (5). In control analyses (see Methods), experiments with CompCor (19) instead of global signal regression yielded similar factors (see Supplemental Results).

The data were band-pass filtered (0.009 Hz ≤ f ≤ 0.08 Hz) and projected onto FreeSurfer fsaverage6 surface space. The projected data were smoothed using a 6mm full-width half-maximum (FWHM) kernel and down-sampled onto FreeSurfer fsaverage5 surface space.

**S3. Resting-state functional connectivity (RSFC) and z-normalization**

We utilized a cortical parcellation (20) consisting of 400 cortical regions-of-interest (ROIs; Figure 1C) as well as a subcortical segmentation (21) consisting of 19 subcortical ROIs (Figure 1D). Specifically, RSFC (Pearson's correlation) was computed among the average time series of 419 brain ROIs (ignoring censored frames), yielding a 419 × 419 RSFC matrix for each participant. Possible effects of age, sex, head motion (mean FD) and site differences were regressed out from all participants' RSFC data with a general linear model (GLM). GENDAAR was treated as a single site because data collection in GENDAAR has been harmonized across sites. Regression coefficients were estimated only from NT participants to retain any ASD-specific interactions with participants' characteristics (e.g., age). The 87571 unique correlation values in each ASD participant's 419 × 419 RSFC matrix were z-normalized with respect to the 348 ABIDE-II+GENDAAR NT participants. A z-score larger (or smaller) than zero for a given ROI pair would imply hyper-connectivity (or hypo-connectivity) relative to NT participants.

When exploring the drawbacks of latent factors for traditional case-control analysis in ABIDE-I, the factor compositions of ABIDE-I ASD participants were inferred using the model parameters estimated from ABIDE-II+GENDAAR. Therefore, the z-normalization was performed with respect to ABIDE-II+GENDAAR NT participants.

**S4. Bayesian model**

We have previously utilized the hierarchical Bayesian model, latent dirichlet allocation (22) (LDA), to estimate latent atrophy factors based on the voxel-based morphometry (VBM) of the structural MRI of Alzheimer's disease (AD) dementia participants (23). LDA was developed to discover topics in a corpus of text documents (22). The idea is that a text document is associated with a distribution of topics (Pr(topic | document)), which are in turn associated with a distribution of words (Pr(word | topic)). LDA is useful because it allows a document to be associated with multiple topics (which can be shared across documents) and each topic to be associated with multiple words (which can be shared across topics).

In the case of AD dementia, we can think of the AD participants as text documents, atrophy factors as topics, and MNI152 voxels as dictionary words. Therefore, each AD participant expresses one or more latent factors, associated with distinct patterns of brain atrophy. When mapping LDA to this work, we can think of ASD participants as text documents, latent ASD factors as topics and ROI pairs as dictionary words. The premise is that each ASD participant expresses one or more latent factors, and each latent factor is associated with distinct but possibly overlapping patterns of hypo/hyper RSFC (Figure 1A).

However, there is one important difference between applying LDA to the VBM data of AD dementia participants and the RSFC data of ASD participants. In the case of VBM, a positive value at a voxel (after z-normalization with respect to controls) indicated atrophy, while a negative value indicated hypertrophy. Since AD dementia is characterized by brain atrophy, we simply set any negative values to zero (23). However, ASD is associated with both hypo-connectivity and hyper-connectivity, so we cannot simply ignore the negative values (hypo-connectivity). As such, we extended the LDA model, so that for each ROI pair, an additional binary variable indicated whether there was hypo-connectivity or hyper-connectivity for the ROI pair.

The precise mathematical formulation is as follows. Let $M$ denote the total number of ASD participants, $K$ denote the number of latent factors, $V$ be the total number of ROI pairs given by the parcellation (i.e., 419 × 418 / 2 = 87571 in our study) and $N_m$ be the number of involved ROI pairs for the $m$-th participant. The model assumes the following generative process. For the $m$-th ($m \in \{1, \dots, M\}$) participant with $K$ factors and $V$ ROI pairs,

1. Randomly draw a mixture of latent factors $\theta \sim$ Dirichlet($\alpha$), where $\alpha$ is the Dirichlet parameter. $\theta$ is a $K$-dimensional vector, where $\sum_k \theta_k = 1$;
2. For the $n$-th ($n \in \{1, \dots, N_m\}$) involved ROI pair $w_n$ of the participant, independently
   (a) Randomly draw a latent factor $z_n \sim$ Multinomial($\theta$);
   (b) Randomly draw an ROI pair $w_n \sim p(w_n \mid z_n, \beta)$. $\beta$ is a $K \times V$ matrix, each row of which defines a multinomial distribution over all the ROI pairs. $\beta_{ij}$ is the probability that the $j$-th ROI pair will be involved in the $i$-th factor;
   (c) Randomly draw an involvement type (i.e., hyper- or hypo-connectivity) for this ROI pair $y_n \sim p(y_n \mid w_n, z_n, \rho)$. $y_n \in \{0, 1\}$, where 1 represents hyper-connectivity, and 0 represents hypo-connectivity. $\rho$ is a $K \times V$ matrix, where each element defines a Bernoulli distribution: $\rho_{ij}$ is the probability that the $j$-th ROI pair will exhibit hyper-connectivity given the $i$-th factor.

The probability that a participant will express the $k$-th latent factor (i.e., Pr(Factor | Participant)) is given by $\theta_k$ ($k \in \{1, \dots, K\}$). The expected factor-specific hypo/hyper RSFC patterns (i.e., E(RSFC patterns | Factor)) is given by $\beta(2\rho - 1)$. For example, for the $j$-th ROI pair and the $i$-th factor, if $\rho_{ij} > 0.5$ (i.e., the $j$-th ROI pair is more likely to exhibit hyper-connectivity given the $i$-th factor), $\beta_{ij}(2\rho_{ij} - 1)$ will be positive, corresponding to hyper-connectivity pattern (hot color in Figure 2A). On the other hand, if $\rho_{ij} < 0.5$ (i.e., the $j$-th ROI pair is more likely to have hypo-connectivity given the $i$-th factor), $\beta_{ij}(2\rho_{ij} - 1)$ will be negative, corresponding to hypo-connectivity pattern (cold color in Figure 2A). If $\beta_{ij}$ is close to 0 (i.e., the $j$-th ROI pair has a low probability of being involved in the $i$-th factor) or if $\rho_{ij}$ is close to 0.5 (i.e., the $j$-th ROI pair has weak preference towards hyper- or hypo-connectivity given the $i$-th factor), then $\beta_{ij}(2\rho_{ij} - 1)$ will be close to 0 (dark color in Figure 2A).

Because word counts for each dictionary word are integers, to apply the Bayesian model to the z-normalized RSFC data from the ASD participants, the z-scores were multiplied by 10 and rounded to the nearest integer. Thus, more positive (negative) values indicated greater hyper-connectivity (hypo-connectivity). While discretization of the RSFC data might lead to some loss of information, we note that for sufficiently large multiplicative factor (10 in this study), there is essentially no loss of information. In practice, we find that this approach works very well (23,24).

**S5. Estimating the Bayesian model**

Given the z-normalized, discretized RSFC data of ASD participants (see previous sections) and a pre-defined number of latent factors $K$, we seek to estimate the probability that an ASD participant is associated with a factor $\theta$, the probability that an ROI pair is associated with a factor $\beta$, and the probability that an ROI pair is hyper-connected $\rho$. Like LDA (22), we used a variational expectation-maximization (VEM) algorithm to estimate these probabilities.

The algorithm iterates between the variational E-step and M-step until convergence. In the E-step, $(\alpha, \beta, \rho)$ are assumed fixed, and the following two equations are iterated until convergence:

$$\phi_{mnk} \propto \exp\left(\Psi(\gamma_{mk}) - \Psi\left(\sum_{i=1}^{K} \gamma_{mi}\right)\right) \beta_{kv(n_m)} \rho_{kv(n_m)}^{y_{mn}} \left(1 - \rho_{kv(n_m)}\right)^{1-y_{mn}}, \quad (1)$$

$$\gamma_{mk} = \alpha + \sum_{n=1}^{N_m} \phi_{mnk}, \quad (2)$$

where $\phi$ is the variational parameter for $z$, so we can think of $\phi_{mnk}$ as the posterior probability that the $n$-th ROI pair of the $m$-th ASD participant is associated with the $k$-th latent factor. $\gamma$ is the variational parameter for $\theta$, so we can think of $\gamma_{mk}$ as the posterior probability that the $m$-th ASD participant is associated with the $k$-th latent factor. $y_{mn}$ is the binary variable indicating whether the $n$-th ROI pair of the $m$-th ASD participant is hyper-connected. $v(n_m)$ indexes the ROI pair (among the 87571 ROI pairs) that the $n$-th ROI pair of the $m$-th ASD participant corresponded to. $\Psi(\cdot)$ is the digamma function (the first derivative of the log Gamma function). $\alpha$ is the Dirichlet parameter.

In the M-step, $\phi$ and $\gamma$ are assumed to be fixed. The hyperparameter $\alpha$ is updated using the Newton-Raphson algorithm (22) and $(\beta, \rho)$ are updated as follows:

$$\beta_{kv} \propto \sum_{m=1}^{M} \sum_{n=1}^{N_m} \phi_{mnk} I(m, n, v), \quad (3)$$

$$\rho_{kv} = \frac{\sum_{m=1}^{M} \sum_{n=1}^{N_m} \phi_{mnk} I(m, n, v) y_{mn}}{\sum_{m=1}^{M} \sum_{n=1}^{N_m} \phi_{mnk} I(m, n, v)}, \quad (4)$$

where $\beta_{kv}$ is the probability that the $v$-th ROI pair (out of 87571 ROI pairs) is associated with the $k$-th factor, $\rho_{kv}$ is the probability that the $v$-th ROI pair (out of 87571 ROI pairs) is associated with hyper-connectivity in the $k$-th factor, $\phi_{mnk}$ is the posterior probability that the $n$-th ROI pair of the $m$-th ASD participant is associated with the $k$-th latent factor, and $y_{mn}$ is the binary variable indicating whether the $n$-th ROI pair of the $m$-th ASD participant is hyper-connected. $\mathrm{I}(m, n, v)$ is a binary variable indicating whether the $n$-th ROI pair of the $m$-th participant corresponds to the $v$-th ROI pair (among the 87571 ROI pairs).

For each predefined number of latent factors $K$, the estimation procedure was repeated with 100 random initializations, resulting in 100 estimates. The final estimate was obtained by selecting the solution closest to the remaining 99 estimates. Briefly, for each estimate, we reordered the latent factors (using the Hungarian matching algorithm) to maximize the correlations of E(RSFC patterns | Factor) between corresponding pairs of latent factors. After obtaining the optimal match, the pairwise correlations were averaged across all latent factors, resulting in an average correlation between each pair of estimates. The estimate having the highest average correlation with the remaining 99 estimates was taken as the final estimate.

#### S6. Comparing participants' characteristics across latent factors by GLM (or logistic regression)

We explored how participants' characteristics (age, sex, FIQ and head motion) varied across the three latent factors using GLM (and logistic regression for sex) in ABIDE-II+GENDAAR. For age and head motion, we used the entire ABIDE-II+GENDAAR ASD cohort of 306 participants. For IQ, there were 9 participants with missing data, so the analysis only involved 297 participants. In the case of sex, only sites with ≥5 female ASD participants (i.e., GU_1, OHSU_1, IP_1, KKI_1 and GENDAAR) were included in the sex analysis. For each site, male participants were selected to match the number, age, FIQ and head motion of female participants, resulting in 114 participants for this analysis.

GLM was applied to compare age, FIQ and head motion across latent factors. The characteristic of interest was treated as response $y$. The probability of a participant belonging to factor 1 ($p_1$) and probability of a participant belonging to factor 2 ($p_2$) were treated as explanatory variables. Nuisance variables consisted of binary indicators of acquisition sites. There were 14 acquisition sites (i.e., 13 sites in ABIDE-II together with GENDAAR), which resulted in 13 nuisance variables: $x_{s1}, x_{s2}, \ldots, x_{s13}$, where each $x$ was a vector of length $M$ (number of ASD participants). The $m$-th element of the $i$-th vector was 1 if the $m$-th participant belonged to the $i$-th acquisition site. Otherwise the value was 0. Thus, the GLM was $y = \beta_0 + \beta_1 p_1 + \beta_2 p_2 + \beta_{s1} x_{s1} + \cdots + \beta_{s13} x_{s13} + \epsilon$, where $\beta$'s were the regression coefficients, and $\epsilon$ was the residual. The probability of factor 3 ($p_3$) was implicitly modeled because $p_1 + p_2 + p_3 = 1$. Intuitively, $\beta_0$ reflected the response of factor 3, $\beta_1$ reflected the response difference between factors 1 and 3, and $\beta_2$ reflected the response difference between factors 2 and 3.

Statistical tests of whether the characteristic $y$ varied across factors involved null hypotheses of the form $H\beta = 0$, where $\beta = [\beta_0, \beta_1, \beta_2, \beta_{s1}, \beta_{s2}, \ldots, \beta_{s13}]^T$, and $H$ is the linear contrast (25). First, we performed a statistical test of overall differences across all latent factors with $H = [0, 1, 0, \ldots, 0; 0, 0, 1, 0, \ldots, 0]$. We then tested for pairwise differences between the factors. For example, $H = [0, -1, 1, 0, \ldots, 0]$ tested possible differences between factors 1 and 2, $H = [0, 1, 0, 0, \ldots, 0]$ compared factors 1 and 3, and $H = [0, 0, 1, 0, \ldots, 0]$ compared factors 2 and 3.

Since sex is a binary variable, logistic regression was used. Here, the response $y$ was sex (zero for male, and one for female). Explanatory variables consisted of the probability of factor 1 ($p_1$) and probability of factor 2 ($p_2$). Like before, nuisance variables consisted of binary indicators of acquisition sites $x_{s1}, \ldots, x_{s4}$ (only 5 sites with ≥5 female ASD participants were included in the sex analysis, i.e., GU_1, OHSU_1, IP_1, KKI_1 and GENDAAR). Thus, the regression model was $\log\left(\frac{\mu}{1-\mu}\right) = \beta_0 + \beta_1 p_1 + \beta_2 p_2 + \beta_{s1} x_{s1} + \cdots + \beta_{s4} x_{s4} + \epsilon$, where $\mu$ was the probability of female, $\beta$'s were the regression coefficients, and $\epsilon$ was the residual. Intuitively, $\exp(\beta_0)$ reflected the odds ratio for factor 3, $\exp(\beta_1)$ reflected the odds ratio between factors 1 and 3, and $\exp(\beta_2)$ reflected the odds ratio between factors 2 and 3.

The likelihood ratio test (25) was used to determine whether sex varied across latent factors. First, we performed a statistical test of overall differences across all latent factors. Here, the restricted model $\log\left(\frac{\mu}{1-\mu}\right) = \beta_0 + \beta_{s1} x_{s1} + \cdots + \beta_{s4} x_{s4} + \epsilon$ was fitted to the data, and the resulting likelihood was compared with the likelihood of the original model $\log\left(\frac{\mu}{1-\mu}\right) = \beta_0 + \beta_1 p_1 + \beta_2 p_2 + \beta_{s1} x_{s1} + \cdots + \beta_{s4} x_{s4} + \epsilon$. Next, we tested for possible pairwise differences between factors. For example, to compare factors 1 and 2, the restricted model was $\log\left(\frac{\mu}{1-\mu}\right) = \beta_0 + \beta_1 (p_1 + p_2) + \beta_{s1} x_{s1} + \cdots + \beta_{s4} x_{s4} + \epsilon$ since $\beta_1 = \beta_2$ under the null hypothesis. To compare factors 1 and 3, the restricted model was $\log\left(\frac{\mu}{1-\mu}\right) = \beta_0 + \beta_2 p_2 + \beta_{s1} x_{s1} + \cdots + \beta_{s4} x_{s4} + \epsilon$ since $\beta_1 = 0$ under the null hypothesis. To compare factors 2 and 3, the restricted model became $\log\left(\frac{\mu}{1-\mu}\right) = \beta_0 + \beta_1 p_1 + \beta_{s1} x_{s1} + \cdots + \beta_{s4} x_{s4} + \epsilon$ since $\beta_2 = 0$ under the null hypothesis.

**S7. Canonical correlation analysis (CCA) between factor loadings and behavioral symptoms**

As explained in the main text, the variables indexing behavioral symptoms were divided into five groups in order to maximize the number of participants in each group. The five groups corresponded to "autistic traits (ADOS)", "restricted/repetitive behaviors", "social responsiveness" and "executive function" and "comorbid psychopathology" (Table S4). Let us denote the factor composition of a participant as $x_1$, $x_2$ and $x_3$ (correspond to Pr(Factor | Participant) after regressing out age, sex, head motion and sites) respectively.

We then performed CCA between each factor loading and each group of behavioral symptoms (also after regressing out age, sex, head motion and sites), i.e., 15 CCAs in total for the three-factor model. For the sake of illustration, let us consider the first factor and the third set of behavioral scores (social responsiveness), consisting of four behavioral scores ("social cognition", "social awareness", "social communication" and "social motivation"). Thus, each participant is associated with a probability of belonging to factor 1 ($x_1$) and four social responsiveness scores. We then performed CCA between $x_1$ and the four behavioral scores. This yielded a maximum of one canonical component because there was only one factor loading score. By construction, this canonical component yielded the optimal linear combination of the four social responsiveness scores that maximally correlate with $x_1$.

To interpret the relative importance of the four social responsiveness scores in this canonical component, the structural coefficient for each of the four social responsiveness scores (26) was computed. Briefly, the canonical component yielded a linear combination of the four social responsiveness scores, resulting in one overall CCA social responsiveness score per participant. To compute the structural coefficient of the social cognition score, we correlated the social cognition score and the overall CCA social responsiveness score across participants. A high positive correlation would imply that higher social cognition score (worse social cognition) was more strongly associated with factor 1 loading. The resulting structural coefficients were plotted as the y-axes (Importance (r)) in the bar plots in Figure 5.

To test if the CCA was statistical significant, a permutation test using the PALM package (27,28) (https://fsl.fmrib.ox.ac.uk/fsl/fslwiki/PALM) was performed. The permutation procedure accounted for the participants coming from different sites, by restricting the permutations to within each site. In the case of factor 1 and the social responsiveness symptoms, the permutation test yielded a p value of 0.004 (Figure 5B), which survived the FDR correction of $q < 0.05$.

**S8. K-means clustering on z-normalized RSFC data of ASD participants**

As a control analysis, we compared our Bayesian model with traditional clustering algorithm. We applied k-means clustering to the z-normalized RSFC data of the 306 ASD participants in ABIDE-II+GENDAAR. K-means clustering was performed using "correlation" as the distance metric (one minus the sample correlation between points) with 1000 random initializations. The solution with the best cost function was chosen to be the final solution. Because the two-factor and four-factor estimates were unstable (see Results and Supplemental Results), we only ran k-means clustering with three clusters as a comparison to the three-factor estimates. The k-means clustering algorithm yields three 419 x 419 RSFC matrices, obtained by averaging across subjects assigned to the corresponding k-means clusters. We then correlated the k-means cluster centers (RSFC matrices) with the factor-specific RSFC patterns (Figure 2A) to examine if the Bayesian latent factors were similar to the k-means cluster centers. We repeated this analysis using rs-fMRI data preprocessed using GSR or CompCor.

**S9. Logistic regression between k-means clusters and behavioral symptoms**

To compare behavioral associations in k-means clusters with those found in the latent factors, we applied logistic regression between k-means clusters (obtained using rs-fMRI data with GSR) and behavioral scores. This analysis was only run on clusters corresponding to latent factors having significant behavioral associations and the respective groups of behavioral scores (see Results and Figure 5). Thus, only four logistic regressions were constructed to investigate associations between cluster 1 and restricted/repetitive behavior, cluster 1 and social responsiveness, cluster 2 and comorbid psychopathology as well as cluster 2 and executive function. The cluster membership was represented by a one-hot vector (1 for belonging to and 0 for not belonging to that cluster) and was treated as response variable $y$. The explanatory variables consisted of z-normalized behavioral scores $s_1, s_2, \dots, s_n$ (where $n$ is the number of behavioral scores). Nuisance variables included age $x_1$, sex $x_2$, head motion $x_3$ (mean FD) and binary indicators of acquisition sites $x_{s1}, \dots, x_{s13}$. Thus, the logistic regression model was $\log\left(\frac{\mu}{1-\mu}\right) = \beta_0 + \beta_1 s_1 + \cdots + \beta_n s_n + \beta_{x1} x_1 + \beta_{x2} x_2 + \beta_{x3} x_3 + \beta_{s1} x_{s1} \dots + \beta_{s13} x_{s13} + \epsilon$, where $\mu$ was the probability of belonging to the cluster, $\beta$'s were the regression coefficients, and $\epsilon$ was the residual. Finally, statistical significance of the logistic regression model was tested using likelihood ratio test (25). Here, the restricted model was $\log\left(\frac{\mu}{1-\mu}\right) = \beta_0 + \beta_{x1} x_1 + \cdots + \beta_{s13} x_{s13} + \epsilon$ because $\beta_1 = \beta_2 = \cdots = \beta_n = 0$ under the null hypothesis. The likelihood of the restricted model was compared with the likelihood of the original model $\log\left(\frac{\mu}{1-\mu}\right) = \beta_0 + \beta_1 s_1 + \cdots + \beta_n s_n + \beta_{x1} x_1 + \beta_{x2} x_2 + \beta_{x3} x_3 + \beta_{s1} x_{s1} \ldots + \beta_{s13} x_{s13} + \epsilon$ to test the statistical significance of the model.

**S10. Relevance of latent factors for traditional case-control analyses in ABIDE-I**

In most studies, ASD and NT individuals are compared without accounting for ASD heterogeneity. To explore the drawbacks of case-control analyses in light of the discovered latent factors from ABIDE-II+GENDAAR, we considered 166 ASD and 150 NT participants from ABIDE-I (Table S1).

First, we computed differences between the 419 x 419 RSFC matrices of the ASD and NT participants. Network-based statistics (29) (NBS) was utilized to correct for multiple comparisons. Second, the factor compositions of ABIDE-I ASD participants were inferred using the hypo/hyper RSFC patterns previously estimated from ABIDE-II+GENDAAR. To allow comparisons with the case-control analysis, ASD participants were assigned to three groups representing the three factors our model allowed each participant to express. We note that assigning an ASD individual to a single group violates the spirit of our approach, but this is necessary for comparison with the traditional case-control analysis.

Here, we experimented with two criteria of subgrouping. (A) If a participant's highest probability Pr(Factor | Participant) was greater than 50%, he/she would be assigned to the corresponding factor. This resulted in 62 ASD participants assigned to the first factor, 17 ASD participants assigned to the second factor, and 10 ASD participants assigned to the third factor. The remaining 77 ASD participants did not have any factor loading greater than 50%, so were not assigned to any factor and were excluded from further analyses. (B) The ASD participants were assigned to the factor with highest probability Pr(Factor | Participant). This resulted in 100 ASD participants assigned to the first factor, 38 ASD participants assigned to the second factor, and 28 ASD participants assigned to the third factor. Note that in this subgrouping criterion, each ASD participant was assigned to a single factor. Finally, RSFC between the ASD and NT participants (with matched age, sex, head motion and sites; see Table S6 and S7) from each of the three groups were compared.

**S11. False discovery rate multiple comparisons correction**

Since multiple statistical tests (bootstrapping, participants characteristics analyses, behavioral CCAs for factors, behavioral logistic regressions for k-means clusters, network-based statistics and t-tests/chi-squared tests) were performed in this study, all p values were corrected using false discovery rate (FDR) at $q = 0.05$ (30).

**S12. Code and data availability**

Code for this work is publicly available at GITHUB_LINK_TO_BE_ADDED. More specifically, the datasets were pre-processed using an in-house pipeline (https://github.com/ThomasYeoLab/CBIG/tree/master/stable_projects/preprocessing/CBIG_fMRI_Preproc2016). Factor compositions of ASD participants and code used in this manuscript are publicly available at GITHUB_LINK_TO_BE_ADDED.

### Supplemental Results

**S1. Control analyses**

We performed several control analyses to examine the robustness of results (see Methods and Supplemental Methods).

First, correlations between original latent factors and factors obtained with CompCor RSFC data are reported in Table S8. Notably, factor 1 and factor 2 from the three-factor model were robust across different preprocessing methods, while factor 3 was inconsistent between preprocessing methods. In addition, the two-factor model produced inconsistent results, thus we focused subsequent control analyses on the three-factor estimates (Table S8).

Next, correlations between the k-means cluster centers and original latent factors are reported in Table S9. K-means clusters obtained from GSR and CompCor RSFC data were similar to the original latent factors, suggesting that our Bayesian model is consistent with the k-means algorithm. Intriguing, factor 3 (GSR) was much more similar to k-means cluster 3 (CompCor) than factor 3 (CompCor).

Thus, overall, factors 1 and 2, and to some extent factor 3, were relatively robust to preprocessing and analysis strategies.

Finally, Figure S4 shows behavioral associations in k-means clusters remaining significant after FDR correction ($q < 0.05$). Importantly, the associations (Figure S4) were in the same direction as the latent factors (Figure 5). However, compared to the latent factors, k-means clusters did not show significant associations with "social responsiveness" or "executive function". This analysis suggests that latent factors given by our Bayesian model exhibit stronger behavioral associations than clusters obtained by traditional k-means clustering algorithm.

Finally, factors estimated from two random splits were highly correlated with the original factors (Table S10A), and were also similar to each other (Table S10B), demonstrating the robustness of the factors.

**S2. Traditional case-control analysis yields smaller effects and misses significant ASD-related RSFC associations**

To explore the relevance of the latent factors for traditional case-control analyses ignoring ASD heterogeneity, we compared the RSFC between all the 166 ASD and 150 NT participants, as well as ASD and NT participants assigned to each ASD subgroup in ABIDE-I (see Methods and Supplemental Methods). We detail the results based on the first subgrouping criterion (i.e., ASD participants were assigned to the factor if Pr(Factor | Participant) > 0.5) (Figure S5). Similar results were obtained using the second subgrouping criterion (Figure S6).

Figure S5A shows the RSFC differences between 166 ASD and 150 NT participants from ABIDE-I. Differences were statistically significant (NBS $p < 0.0001$). The patterns of hypo/hyper RSFC were similar to previous results from ABIDE-I (1,31). Figures S5B shows the underlying factor composition of the ABIDE-I participants (inferred with model parameters estimated from ABIDE-II+GENDAAR). Each dot represents a participant. Red, blue and green dots indicate participants assigned to factors 1, 2 and 3 respectively. Gray dots represent participants not assigned to any group. The three ASD sub-groups were in turn compared with matched NT participants separately (Figure S5C) yielding statistically significant results for all three contrasts (NBS $p < 0.0001$).

Importantly, RSFC differences for factor 1 participants (Figure S5C) were most similar to traditional case-control analyses (Figure S5A; $r = 0.56$, 0.14 and 0.17 for factor 1, 2 and 3 respectively) because factor 1 was most strongly expressed among ABIDE-I ASD participants (Figure S5B). Consequently, RSFC differences associated with factors 2 and 3 (Figure S5C) were not detected in the traditional case-control analysis (Figure S5A). Finally, when comparing magnitudes (effect sizes) in Figures S5A and S5C, it is obvious that traditional case-control analysis (despite the larger sample size) yielded significantly weaker RSFC differences than the subgroup analyses.

#### Supplemental Tables and Figures

**Table S1. Characteristics of participants from ABIDE-I.** After quality control (Supplemental Methods), eight data collections from ABIDE-I were included in this study: TRINITY ($N_{ASD}$ = 18, $N_{NT}$ = 21), USM ($N_{ASD}$ = 21, $N_{NT}$ = 21), YALE ($N_{ASD}$ = 16, $N_{NT}$ = 13), LEUVEN_1 ($N_{ASD}$ = 10, $N_{NT}$ = 10), KKI ($N_{ASD}$ = 8, $N_{NT}$ = 16), NYU ($N_{ASD}$ = 54, $N_{NT}$ = 40), UCLA_1 ($N_{ASD}$ = 30, $N_{NT}$ = 20), UCLA_2 ($N_{ASD}$ = 9, $N_{NT}$ = 9). ASD and NT participants were compared with either two-sample t-tests (for continuous measures) or chi-squared tests (for categorical measures). All p-values that survived false discovery rate (FDR) correction (q < 0.05) are indicated in bold.

| | ***ASD* (N = 166)** | ***NT* (N = 150)** | ***p value* (two-tailed)** |
|---|---|---|---|
| *Demographics* | | | |
| Age, mean (SD), years | 15.12 (6.19) | 15.98 (6.08) | 0.22 |
| Female, No. (%) | 19 (11.45) | 24 (16.00) | 0.31 |
| Full-scale IQ, mean (SD) | 103.56 (15.80) | 110.16 (12.99) | **6.97e-5** |
| *Head motion* | | | |
| Mean FD before censoring, mean (SD) | 0.08 (0.04) | 0.07 (0.03) | 0.12 |
| Mean FD after censoring, mean (SD) | 0.05 (0.01) | 0.05 (0.01) | 0.91 |
| *Current medication (by target)*[a] | | | |
| GABA, No. (%) | 1 (0.78) | 0 (0.00) | 0.96 |
| Glutamate, No. (%) | 2 (1.55) | 0 (0.00) | 0.53 |
| Serotonin, No. (%) | 21(16.28) | 0 (0.00) | **1.59e-5** |
| Epinephrine/Adrenaline, No. (%) | 0 (0.00) | 0 (0.00) | NA |
| Norepinephrine/Noradrenaline, No. (%) | 23 (17.83) | 1 (0.86) | **2.18e-5** |
| Dopamine, No. (%) | 21(16.28) | 1 (0.86) | **6.59e-5** |
| Others, No. (%) | 1(0.78) | 1 (0.86) | 0.53 |
| *Current medication (by class)*[b] | | | |
| Antipsychotic, No. (%) | 10 (7.75) | 0 (0.00) | **6.20e-3** |
| Antidepressant, No. (%) | 17 (13.18) | 0 (0.00) | **1.44e-4** |
| Stimulant, No. (%) | 13 (10.08) | 1 (0.86) | **4.70e-3** |
| SHA, No. (%) | 1 (0.78) | 0 (0.00) | 0.96 |
| Mood stabilizer, No. (%) | 2 (1.55) | 0 (0.00) | 0.53 |
| Others, No. (%) | 9 (6.98) | 1 (0.86) | 0.04 |

[a]Medication was sorted by the neurotransmitter system(s) targeted by the medication currently used by participants, based on the Neuroscience-based Nomenclature (NbN-2 (32,33); http://www.nbn2.com/). Only 129 ASD and 116 NT participants from ABIDE-I had medication information. The percentages were computed based on 129 ASD and 116 NT participants.
[b]Only 129 ASD and 116 NT participants from ABIDE-I had medication information. The percentages were computed based on 129 ASD and 116 NT participants.
*Abbreviations*: GABA, Gamma-Aminobutyric acid; SHA, sedatives/hypnotics/anxiolytics.

**Table S2. Number of participants excluded in censoring step in fMRI pre-processing.** In fMRI pre-processing censoring step (see Supplemental Methods), functional runs with more than half of the volumes labeled as censored frames were completely removed, resulting in some participants being excluded. The participants in each site that were excluded in the censoring step are shown in the table below.

| Repositories | Number of ASD participants excluded | Number of NT participants excluded |
|---|---|---|
| **ABIDE-I** | $N_{TRINITY} = 2$<br>$N_{USM} = 5$<br>$N_{YALE} = 3$<br>$N_{KKI} = 11$<br>$N_{NYU} = 3$<br>$N_{UCLA\_1} = 9$<br>$N_{UCLA\_2} = 3$ | $N_{TRINITY} = 2$<br>$N_{USM} = 4$<br>$N_{YALE} = 2$<br>$N_{LEUVEN\_1} = 1$<br>$N_{KKI} = 7$<br>$N_{NYU} = 3$<br>$N_{UCLA\_1} = 3$ |
| **ABIDE-II** | $N_{OILH\_2} = 2$<br>$N_{GU\_1} = 12$<br>$N_{SDSU\_1} = 1$<br>$N_{BNI\_1} = 7$<br>$N_{ETH\_1} = 6$<br>$N_{TCD\_1} = 7$<br>$N_{NYU\_1} = 4$<br>$N_{KKI\_1} = 25$<br>$N_{USM\_1} = 7$<br>$N_{IU\_1} = 2$<br>$N_{IP\_1} = 2$<br>$N_{KUL\_3} = 1$<br>$N_{UCLA\_1} = 5$<br>$N_{EMC\_1} = 10$<br>$N_{UCD\_1} = 2$<br>$N_{U\_MIA\_1} = 1$ | $N_{OILH\_2} = 1$<br>$N_{GU\_1} = 6$<br>$N_{SDSU\_1} = 1$<br>$N_{BNI\_1} = 14$<br>$N_{ETH\_1} = 2$<br>$N_{TCD\_1} = 3$<br>$N_{NYU\_1} = 1$<br>$N_{KKI\_1} = 42$<br>$N_{USM\_1} = 1$<br>$N_{IP\_1} = 4$<br>$N_{UCLA\_1} = 4$<br>$N_{EMC\_1} = 10$<br>$N_{U\_MIA\_1} = 2$ |
| **GENDAAR** | $N_{ASD} = 43$ | $N_{NT} = 34$ |

**Table S3. Behavioral data of 306 ASD participants from ABIDE-II+GENDAAR.** For each behavioral measure, only a subset of ASD participants have the scores available.

| Scale | Subscale/Measure | Mean (SD) |
|---|---|---|
| Autism Diagnostic Observation Schedule (ADOS) scores | Social (N = 176) | 7.25 (2.58) |
| | Communication (N = 176) | 3.23 (1.50) |
| | Stereotyped behavior (N = 176) | 1.64 (1.40) |
| Social Responsiveness Scale (SRS) raw scores | Social cognition (N = 200) | 16.62 (6.27) |
| | Social awareness (N = 200) | 11.92 (3.97) |
| | Social communication (N = 200) | 31.40 (11.38) |
| | Social motivation (N = 200) | 15.29 (6.66) |
| | Autistic mannerism (N = 200) | 17.41 (6.91) |
| Repetitive Behaviors Scale-Revised 6 Subscales (RBSR-6) scores | Stereotyped behavior (N = 92) | 3.55 (3.05) |
| | Self-injurious behavior (N = 92) | 1.99 (2.92) |
| | Compulsive behavior (N = 92) | 3.72 (4.24) |
| | Ritualistic behavior (N = 92) | 5.01 (3.91) |
| | Sameness behavior (N = 92) | 7.95 (6.23) |
| | Restricted interests (N = 92) | 3.41 (2.71) |
| Behavior Rating Inventory of Executive Function (BRIEF) T scores | Inhibit (N = 171) | 61.43 (12.64) |
| | Shift (N = 171) | 68.88 (14.17) |
| | Emotional control (N = 171) | 61.43 (12.79) |
| | Initiate (N = 171) | 64.96 (11.26) |
| | Working memory (N = 171) | 66.64 (11.55) |
| | Plan/Organize (N = 171) | 65.70 (12.55) |
| | Organization of materials (N = 171) | 58.51 (10.52) |
| | Monitor (N = 171) | 64.31 (12.27) |
| Child Behavior Checklist Ages 6-18 (CBCL-6-18) scores | Externalizing problems (N = 169) | 55.49 (10.36) |
| | Internalizing problems (N = 169) | 62.99 (10.05) |
| | Affective problems (N = 123) | 63.40 (9.11) |
| | Anxiety problems (N = 123) | 63.26 (8.94) |
| | Somatic problems (N = 123) | 57.70 (8.87) |
| | Attention deficit/hyperactivity problems (N = 123) | 63.10 (8.42) |
| | Oppositional defiant problems (N = 123) | 58.63 (8.67) |
| | Conduct problems (N = 123) | 56.30 (7.90) |

**Table S4. Five groups of behavioral scores used in Canonical Correlation Analysis (CCA) to examine associations between latent factors and behavioral symptoms in ABIDE-II+GENDAAR.** Because ABIDE-II+GENDAAR consisted of datasets across independent sites, participants from different sites might not have the same behavioral scores. If we considered all available behavioral measures jointly, we would be left with only seven participants. Therefore, available behavioral scores were divided into five groups, while maximizing the number of participants in each group. ADOS Stereotyped behavior score was not grouped together in the Restricted/repetitive behaviors domain because only 38 participants have ADOS, RBSR-6 and SRS scores jointly.

| | **Behavioral scores included** |
|---|---|
| **Autistic traits (N = 176[a])** | ADOS Social<br>ADOS Communication<br>ADOS Stereotyped behavior |
| **Restricted/repetitive behaviors (N = 76[b])** | RBSR-6 Stereotyped behavior<br>RBSR-6 Self-injurious behavior<br>RBSR-6 Compulsive behavior<br>RBSR-6 Ritualistic behavior<br>RBSR-6 Sameness behavior<br>RBSR-6 Restricted interests<br>SRS Autistic mannerisms |
| **Social responsiveness (N = 200[c])** | SRS Social awareness<br>SRS Social cognition<br>SRS Social communication<br>SRS Social motivation |
| **Executive function (N = 171[d])** | BRIEF Inhibit<br>BRIEF Shift<br>BRIEF Emotional control<br>BRIEF Initiate<br>BRIEF Working memory<br>BRIEF Plan/Organize<br>BRIEF Organization of materials<br>BRIEF Monitor |
| **Comorbid psychopathology (N = 123[e])** | CBCL-6-18 Internalizing problems<br>CBCL-6-18 Externalizing problems<br>CBCL-6-18 Affective problems<br>CBCL-6-18 Anxiety problems<br>CBCL-6-18 Somatic problems<br>CBCL-6-18 Attention deficit/hyperactivity problems<br>CBCL-6-18 Oppositional defiant problems<br>CBCL-6-18 Conduct problems |

[a]ASD participants from the following sites were included in this analysis: GU_1 (N = 28); OHSU_1 (N = 2); BNI_1 (N = 20); ETH_1 (N = 6); TCD_1 (N = 14); NYU_1 (N = 34); KKI_1 (N = 11); USM_1 (N = 6); IU_1 (N = 17); IP_1 (N = 10); UCLA_1 (N = 9); UCD_1 (N = 1); GENDAAR (N = 18).
[b]ASD participants from the following sites were included in this analysis: BNI_1 (N = 14); TCD_1 (N = 14); NYU_1 (N = 7); KKI_1 (N = 26); UCD_1 (N = 15).

[c]ASD participants from the following sites were included in this analysis: GU_1 (N = 34); OHSU_1 (N = 20); BNI_1 (N = 20); ETH_1 (N = 5); TCD_1 (N = 14); NYU_1 (N = 42); KKI_1 (N = 30); USM_1 (N = 7); IU_1 (N = 13); UCD_1 (N = 15).
[d]ASD participants from the following sites were included in this analysis: GU_1 (N = 34); NYU_1 (N = 39); KKI_1 (N = 30); U_MIA_1 (N = 8); GENDAAR (N = 60).
[e]ASD participants from the following sites were included in this analysis: GU_1 (N = 34); KKI_1 (N = 27); GENDAAR (N = 62).
*Abbreviations*: ADOS, Autism Diagnostic Observation Schedule; RBSR-6, Repetitive Behaviors Scale-Revised 6 Subscales; SRS, Social Responsiveness Scale; BRIEF, Behavior Rating Inventory of Executive Function; CBCL-6-18, Child Behavior Checklist Ages 6-18.

**Table S5. Characteristics of participants in split-half control analysis.** We randomly split all 306 ASD participants from ABIDE-II+GENDAAR into two sets, and matched age, sex, FIQ, head motion (mean FD before censoring) and sites between the two sets. The two sets were compared with either two-sample t-tests (for continuous measures) or chi-squared tests (for categorical measures).

| | **Set 1 ($N_{ASD}$ = 153)** | **Set 2 ($N_{ASD}$ = 153)** | ***p value (two-tailed)*** |
|---|---|---|---|
| *Demographics* | | | |
| Age, mean (SD), years | 14.79 (8.40) | 15.09 (8.80) | 0.76 |
| Female, No. (%) | 29 (18.95) | 41 (26.80) | 0.13 |
| Full-scale IQ, mean (SD)[a] | 106.38 (17.03) | 106.34 (16.83) | 0.98 |
| *Head motion* | | | |
| Mean FD before censoring, mean (SD) | 0.09 (0.05) | 0.09 (0.06) | 0.57 |
| Mean FD after censoring, mean (SD) | 0.05 (0.02) | 0.05 (0.02) | 0.13 |

[a]Only 149 and 148 ASD participants have full-scale IQ scores in set 1 and set 2 respectively.

**Table S6. Characteristics of participants assigned in each factor group (K=3) based on the first subgrouping criterion in the relevance for traditional case-control analyses on ABIDE-I sample.** The ASD participants in ABIDE-I were assigned to three groups: if a participant's highest probability Pr(Factor | Participant) was greater than 50%, he/she would be assigned to the corresponding factor. This resulted in 89 out of 166 ASD participants assigned. In each factor group, the NT participants were matched with ASD participants by age, sex and mean FD (before censoring) within each site. ASD and NT participants were compared with either two-sample t-tests (for continuous measures) or chi-squared tests (for categorical measures).

| | **Factor 1 Group** | | | **Factor 2 Group** | | | **Factor 3 Group** | | |
|---|---|---|---|---|---|---|---|---|---|
| | **ASD (N = 62[a])** | **NT (N = 150[b])** | ***p value (two-tailed)*** | **ASD (N = 17[c])** | **NT (N = 114[d])** | ***p value (two-tailed)*** | **ASD (N = 10[e])** | **NT (N = 115[f])** | ***p value (two-tailed)*** |
| *Demographics* | | | | | | | | | |
| Age, mean (SD), years | 15.08 (7.55) | 15.98 (6.08) | 0.36 | 16.52 (3.42) | 17.14 (6.41) | 0.70 | 18.70 (7.66) | 16.40 (6.03) | 0.26 |
| Female, No. (%) | 6 (9.68) | 24 (16.00) | 0.32 | 1 (5.88) | 14 (12.28) | 0.72 | 2 (20.00) | 16 (13.91) | 0.96 |
| Full-scale IQ, mean (SD) | 105.53 (17.46) | 110.16 (12.99) | 0.03 | 106.24 (19.93) | 110.65 (13.62) | 0.25 | 104.70 (10.37) | 108.75 (13.29) | 0.35 |
| *Head motion* | | | | | | | | | |
| Mean FD before censoring, mean (SD) | 0.08 (0.03) | 0.07 (0.03) | 0.12 | 0.08 (0.07) | 0.07 (0.03) | 0.23 | 0.06 (0.03) | 0.07 (0.03) | 0.38 |
| Mean FD after censoring, mean (SD) | 0.05 (0.01) | 0.05 (0.01) | 0.26 | 0.05 (0.01) | 0.05 (0.01) | 0.52 | 0.04 (0.01) | 0.05 (0.01) | 0.26 |

[a]ASD participants from the following sites were assigned in this group: TRINITY (N = 2); USM (N = 7); YALE (N = 6); LEUVEN_1 (N = 7); KKI (N = 2); NYU (N = 22); UCLA_1 (N = 15); UCLA_2 (N = 1).
[b]All NT participants in ABIDE-I sample were included.
[c]ASD participants from the following sites were assigned in this group: TRINITY (N = 4); USM (N = 3); YALE (N = 3); LEUVEN_1 (N = 1); NYU (N = 4); UCLA_2 (N = 2).
[d]NT participants from the following sites were assigned in this group: TRINITY (N = 21); USM (N = 21); YALE (N = 13); LEUVEN_1 (N = 10); NYU (N = 40); UCLA_2 (N = 9).
[e]ASD participants from the following sites were assigned in this group: TRINITY (N = 1); USM (N = 5); YALE (N = 1); NYU (N = 1); UCLA_1 (N = 2).
[f]NT participants from the following sites were assigned in this group: TRINITY (N = 21); USM (N = 21); YALE (N = 13); NYU (N = 40); UCLA_1 (N = 20).

**Table S7. Characteristics of participants assigned in each factor group (K=3) based on the second subgrouping criterion in the relevance for traditional case-control analyses on ABIDE-I sample.** The ASD participants in ABIDE-I were assigned to the corresponding factor group with highest probability Pr(Factor | Participant). In each factor group, the NT participants were matched with ASD participants by age, sex and mean FD (before censoring) within each site. ASD and NT participants were compared with either two-sample t-tests (for continuous measures) or chi-squared tests (for categorical measures). All p-values that survived false discovery rate (FDR) correction (q < 0.05) are indicated in bold.

| | Factor 1 Group | | | Factor 2 Group | | | Factor 3 Group | | |
|---|---|---|---|---|---|---|---|---|---|
| | **ASD (N = 100[a])** | **NT (N = 150[b])** | **p value (two-tailed)** | **ASD (N = 38[c])** | **NT (N = 150[d])** | **p value (two-tailed)** | **ASD (N = 28[e])** | **NT (N = 140[f])** | **p value (two-tailed)** |
| *Demographics* | | | | | | | | | |
| Age, mean (SD), years | 14.76 (6.53) | 15.98 (6.08) | 0.13 | 15.33 (4.33) | 15.98 (6.08) | 0.54 | 16.14 (7.12) | 15.43 (5.88) | 0.57 |
| Female, No. (%) | 9 (9.00) | 24 (16.00) | 0.16 | 3 (7.89) | 24 (16.00) | 0.31 | 7 (25.00) | 24 (17.14) | 0.48 |
| Full-scale IQ, mean (SD) | 104.56 (15.88) | 110.16 (12.99) | **2.52e-3** | 102.29 (16.84) | 110.16 (12.99) | **2.02e-3** | 101.71 (14.21) | 109.79 (12.89) | **3.38e-3** |
| *Head motion* | | | | | | | | | |
| Mean FD before censoring, mean (SD) | 0.08 (0.04) | 0.07 (0.03) | 0.07 | 0.08 (0.05) | 0.07 (0.03) | 0.30 | 0.07 (0.05) | 0.07 (0.03) | 0.89 |
| Mean FD after censoring, mean (SD) | 0.05 (0.01) | 0.05 (0.01) | 0.07 | 0.05 (0.01) | 0.05 (0.01) | 0.57 | 0.04 (0.01) | 0.05 (0.01) | **3.4e-3** |

[a]ASD participants from the following sites were assigned in this group: TRINITY (N = 10); USM (N = 8); YALE (N = 9); LEUVEN_1 (N = 8); KKI (N = 3); NYU (N = 35); UCLA_1 (N = 22); UCLA_2 (N = 5).
[b]All NT participants in ABIDE-I sample were included.
[c]ASD participants from the following sites were assigned in this group: TRINITY (N = 6); USM (N = 8); YALE (N = 5); LEUVEN_1 (N = 2); KKI (N = 3); NYU (N = 9); UCLA_1 (N = 2); UCLA_2 (N = 3).
[d]All NT participants in ABIDE-I sample were included.
[e]ASD participants from the following sites were assigned in this group: TRINITY (N = 2); USM (N = 5); YALE (N = 2); KKI (N = 2); NYU (N = 10); UCLA_1 (N = 6); UCLA_2 (N = 1).
[f]NT participants from the following sites were assigned in this group: TRINITY (N = 21); USM (N = 21); YALE (N = 13); KKI (N = 16); NYU (N = 40); UCLA_1 (N = 20); UCLA_2 (N = 9).

**Table S8. Correlations between original factors and CompCor factors in control analyses.** Instead of global signal regression (GSR), we used CompCor in the preprocessing step and re-estimated latent factors with two- and three-factor models. Because the two-factor model produced inconsistent results between the two preprocessing methods, we focused on the three-factor model in our manuscript.

| | Factor 1 | Factor 2 | Factor 3 |
|---|---|---|---|
| **CompCor factors, K = 2** | 0.33 | 0.24 | NA |
| **CompCor factors, K = 3** | 0.58 | 0.66 | 0.31 |

**Table S9. Correlations between latent factors and k-means clusters in control analyses with three clusters.** GSR k-means clusters: k-means clusters obtained using RSFC data with global signal regressed. CompCor k-means clusters: k-means clusters obtained using RSFC data preprocessed with CompCor instead of global signal regression.

| | Factor 1 | Factor 2 | Factor 3 |
|---|---|---|---|
| **GSR k-means clusters** | 0.84 | 0.79 | 0.85 |
| **CompCor k-means clusters** | 0.53 | 0.70 | 0.60 |

**Table S10. Correlations across factors in split-half control analysis.** (A) Correlations between latent factors in ABIDE-II+GENDAAR and factors in set 1 and set 2. (B) Correlations between factors in set1 and set 2.

(A)

| | Factor 1 | Factor 2 | Factor 3 |
|---|---|---|---|
| **Set 1 factors** | 0.85 | 0.88 | 0.74 |
| **Set 2 factors** | 0.88 | 0.85 | 0.90 |

(B)

| | Set 1 factor 1 | Set 1 factor 2 | Set 1 factor 3 |
|---|---|---|---|
| **Set 2 factors** | 0.60 | 0.62 | 0.49 |

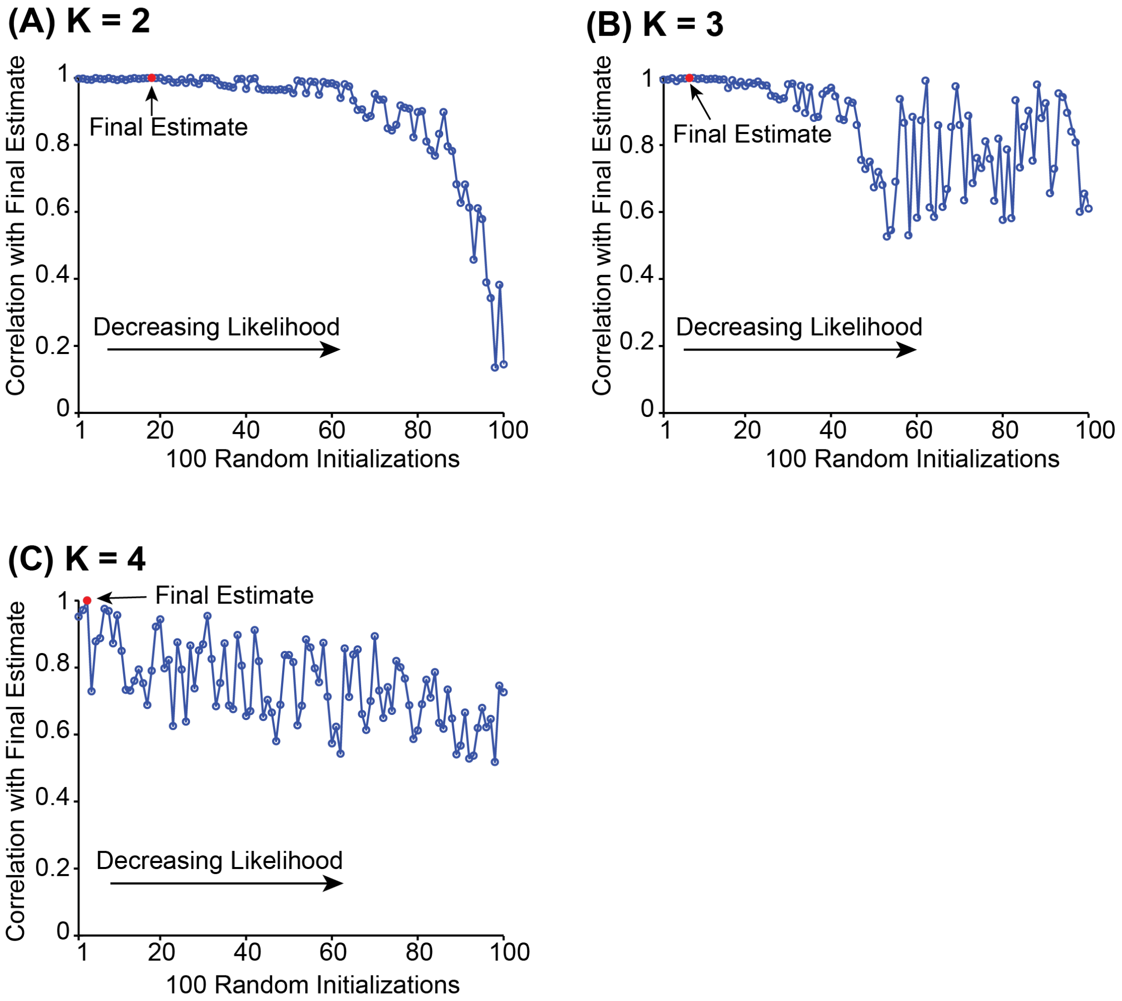


**Figure S1. Correlations between the final estimate and solutions from the 100 random initializations in two-, three-, and four-factor estimates.** This figure shows the correlations between the final estimate (i.e., solution having the highest average correlation with other solutions; see Supplemental Methods) and solutions from the 100 random initializations when variational expectation-maximization (VEM) algorithm was applied to estimate Pr(Factor | Participant) and E(RSFC patterns | Factor) for each number of latent factors K. The solutions were ordered from the highest likelihood to the lowest likelihood. The final estimate is highlighted in red. The 100 random initializations led to robust factor estimations for (A) K = 2 and (B) K = 3, but unstable factor estimations for (C) K = 4. So larger number of latent factors was not considered.

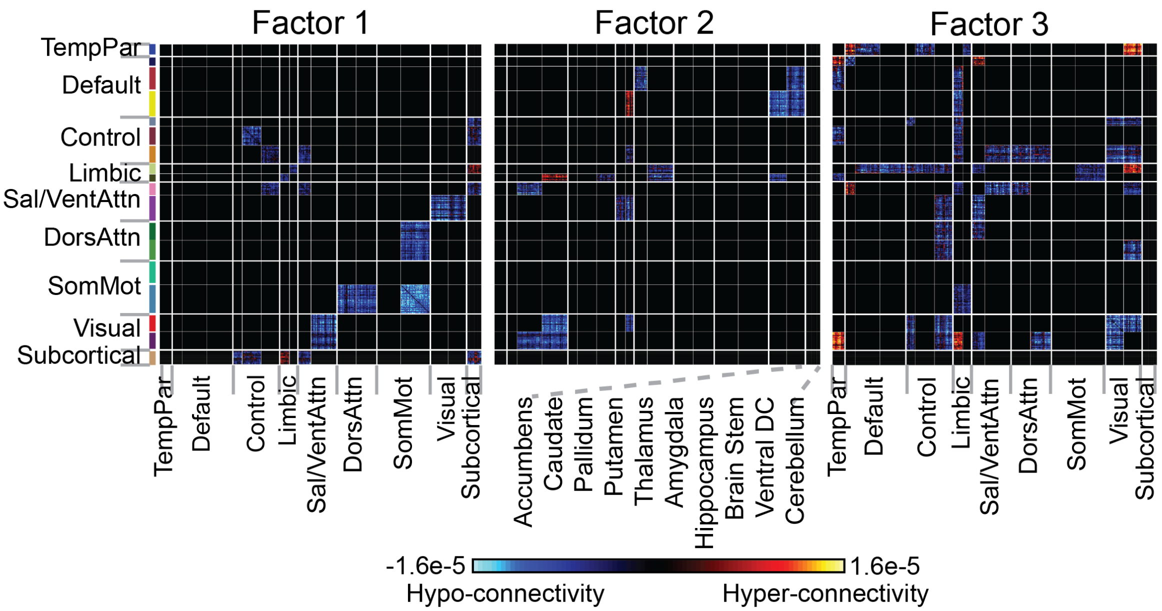


**Figure S2. Unique hypo/hyper RSFC associated with each factor.** Hypo/hyper RSFC unique to each factor was obtained by comparing the three subplots in Figure 2B. Factor 1 was uniquely characterized by hypo-connectivity within somatomotor A, as well as between somatomotor A and dorsal attention, between visual and salience/ventral attention A, as well as hyper-connectivity between limbic B and subcortical regions. Factor 2 was uniquely characterized by hypo-connectivity between default and visual networks, between default B and salience/ventral attention B, as well as hyper-connectivity between limbic A and default A networks. Lastly, Factor 3 was uniquely characterized by a complex hypo/hyper RSFC pattern, such as hypo-connectivity within visual networks and hyper-connectivity between visual A and temp-par, limbic B networks.

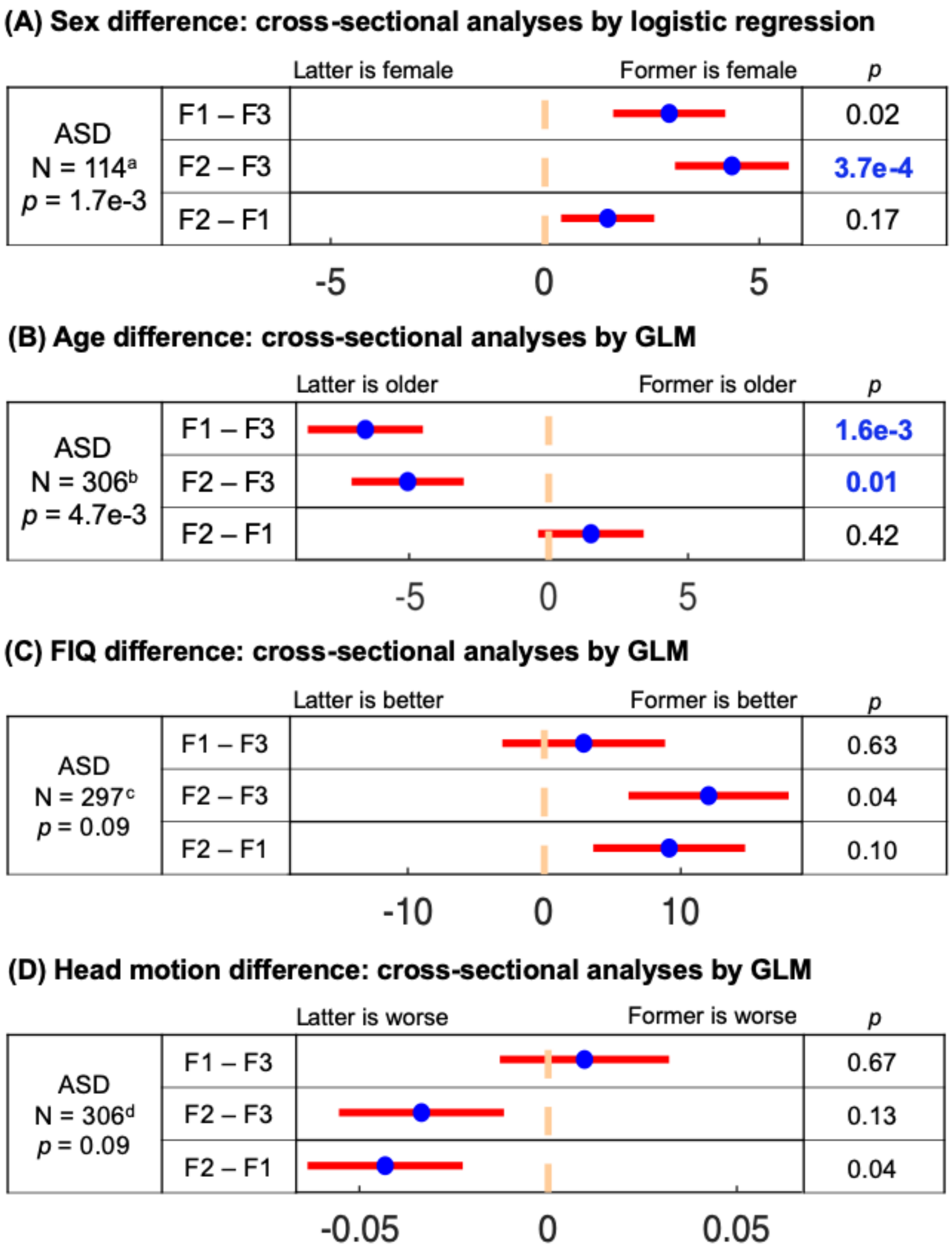


**Figure S3. Third latent factor was associated with males and older participants, and no difference in FIQ or head motion across factors.** Comparisons of (A) sex, (B) age, (C) full-scale IQ and (D) head motion in ASD participants in ABIDE-II+GENDAAR across factors. Pairwise comparisons remaining significant after false discovery rate (FDR; $q < 0.05$) multiple comparisons correction are highlighted in blue. Blue dots are estimated differences between factors, and red bars correspond to standard errors. For example, the second row (F2 – F3) in (A) implies that factor 2 was more strongly associated with females than factor 3.

No pairwise comparison remained significant after false discovery rate (FDR; $q < 0.05$) multiple comparisons correction in (C) and (D), suggesting that latent factors were not simply reflecting FIQ or motion.
*Abbreviations*: F1, factor 1; F2, factor 2; F3, factor 3.

[a]Only sites with ≥5 female ASD participants (i.e., GU_1, OHSU_1, IP_1, KKI_1 and GENDAAR) were included in the sex analysis. For each site, male participants were selected to match the number of female participants, as well as the age, FIQ and head motion of female participants, resulting in 114 participants for this analysis. $N_{GU\_1,male} = N_{GU\_1,female} = 6$; $N_{OHSU\_1,male} = N_{OHSU\_1,female} = 5$; $N_{KKI\_1,male} = N_{KKI\_1,female} = 9$; $N_{IP\_1,male} = N_{IP\_1,female} = 5$; $N_{GENDAAR,male} = N_{GENDAAR,female} = 32$.
[b]All ASD participants in ABIDE-II+GENDAAR were included in this analysis.
[c]Only 297 ASD participants in ABIDE-II+ GENDAAR sample had full-scale IQ scores.
[d]All ASD participants in ABIDE-II+GENDAAR were included.

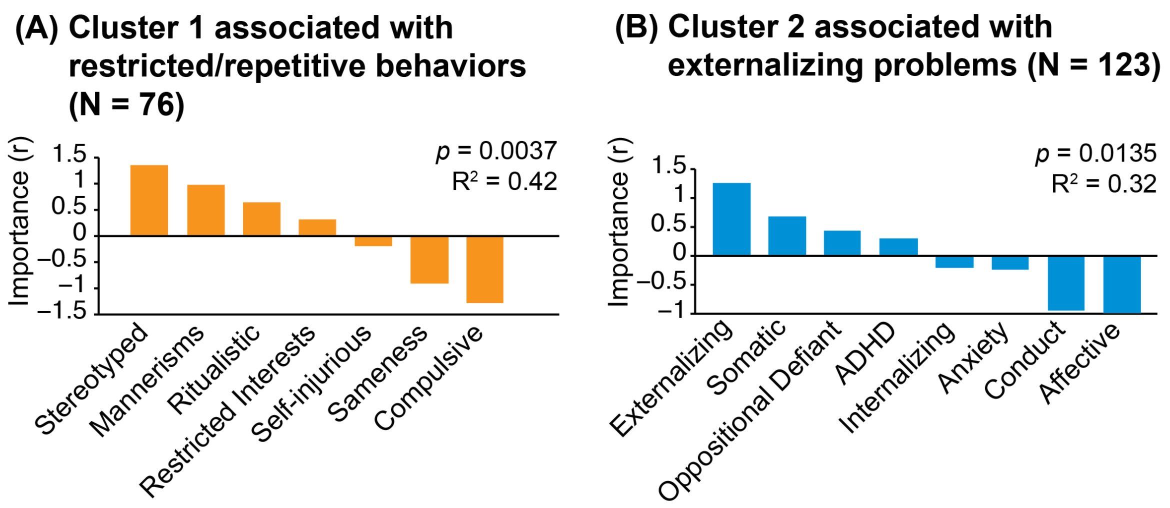


**Figure S4. Behavioral associations in k-means clusters by logistic regression.** Two sets of logistic regression analyses remained significant after FDR ($q < 0.05$) multiple comparisons correction. (A) Associations between cluster 1 and RRB measured by SRS Autistic Mannerism subscale and RBSR-6 subscales. The directions of associations are similar to that in factor 1 CCA analysis (Figure 5A). (B) Associations between cluster 2 and comorbid psychopathology. The directions of associations are similar to that in factor 2 CCA analysis (Figure 5C). Positive correlation suggests that the participants in the cluster had greater impairment. In comparison to the latent factors, k-means clusters did not show significant association with social responsiveness measured by SRS subscales, neither with executive function measured by BRIEF subscales.

*Abbreviations*: SRS, Social Responsiveness Scale; RBSR-6, Repetitive Behaviors Scale-Revised 6 Subscales; CBCL-6-18, Child Behavior Checklist Ages 6-18; BRIEF, Behavior Rating Inventory of Executive Function.

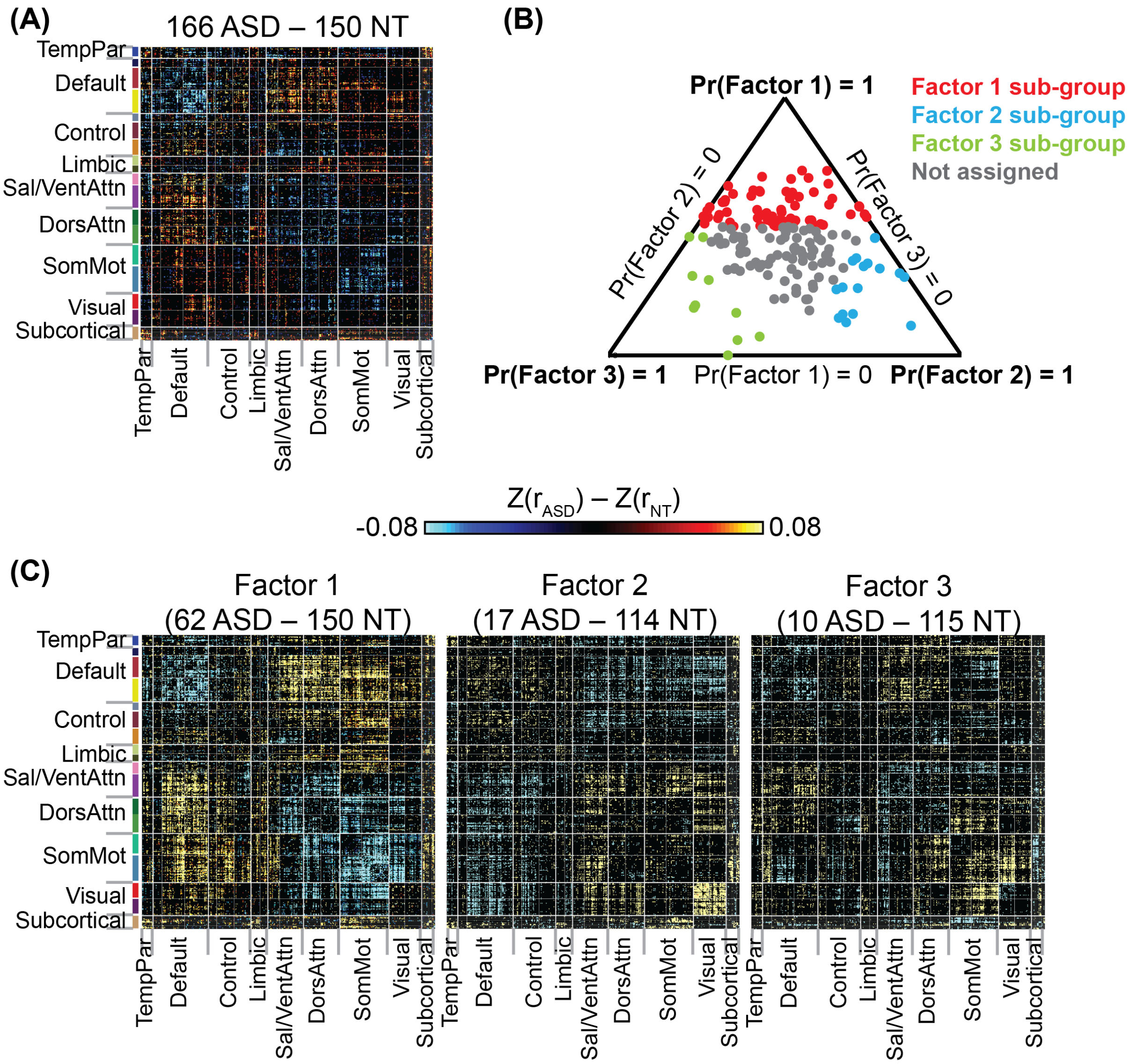


**Figure S5. Traditional case-control analyses (first subgrouping criterion) yielded weaker effects, while missing significant ASD-related RSFC associations.** (A) Traditional case-control analysis: RSFC differences between 166 ASD and 150 NT participants from ABIDE-I matched for age, sex and head motion within each site. ROI pairs with t-statistic > 2 are colored. RSFC differences were statistically significant after correcting for multiple comparisons (NBS $p < 0.0001$). Hot (cold) color indicates hyper-connectivity (hypo-connectivity) in ASD compared to NT participants. (B) Factor compositions of ABIDE-I ASD participants inferred using model parameters estimated from ABIDE-II+GENDAAR. Each participant corresponds to a dot, whose location (in barycentric coordinates) represents the factor composition. Corners of the triangle represent pure factors; closer distance to the respective corner indicates higher probability for the respective factor. Each ASD participant whose highest Pr(Factor | Participant) greater than 50% was assigned to the corresponding factor. Red, blue and green dots represent participants assigned to factor 1 (N = 62), factor 2 (N = 17) and factor 3 (N = 10) respectively. Gray dots represent participants not assigned to any group. (C) RSFC differences between ASD and matched NT participants in the three groups (NBS $p < 0.0001$ for all three analyses). Effects were significantly stronger in sub-group analyses compared with case-control analyses despite the smaller sample sizes. Furthermore, case-control analysis showed similar hypo/hyper RSFC patterns as factor 1 ASD participants ($r = 0.56$), but missed out on RSFC differences shown by participants assigned to factors 2 ($r = 0.14$) and factor 3 ($r = 0.17$).

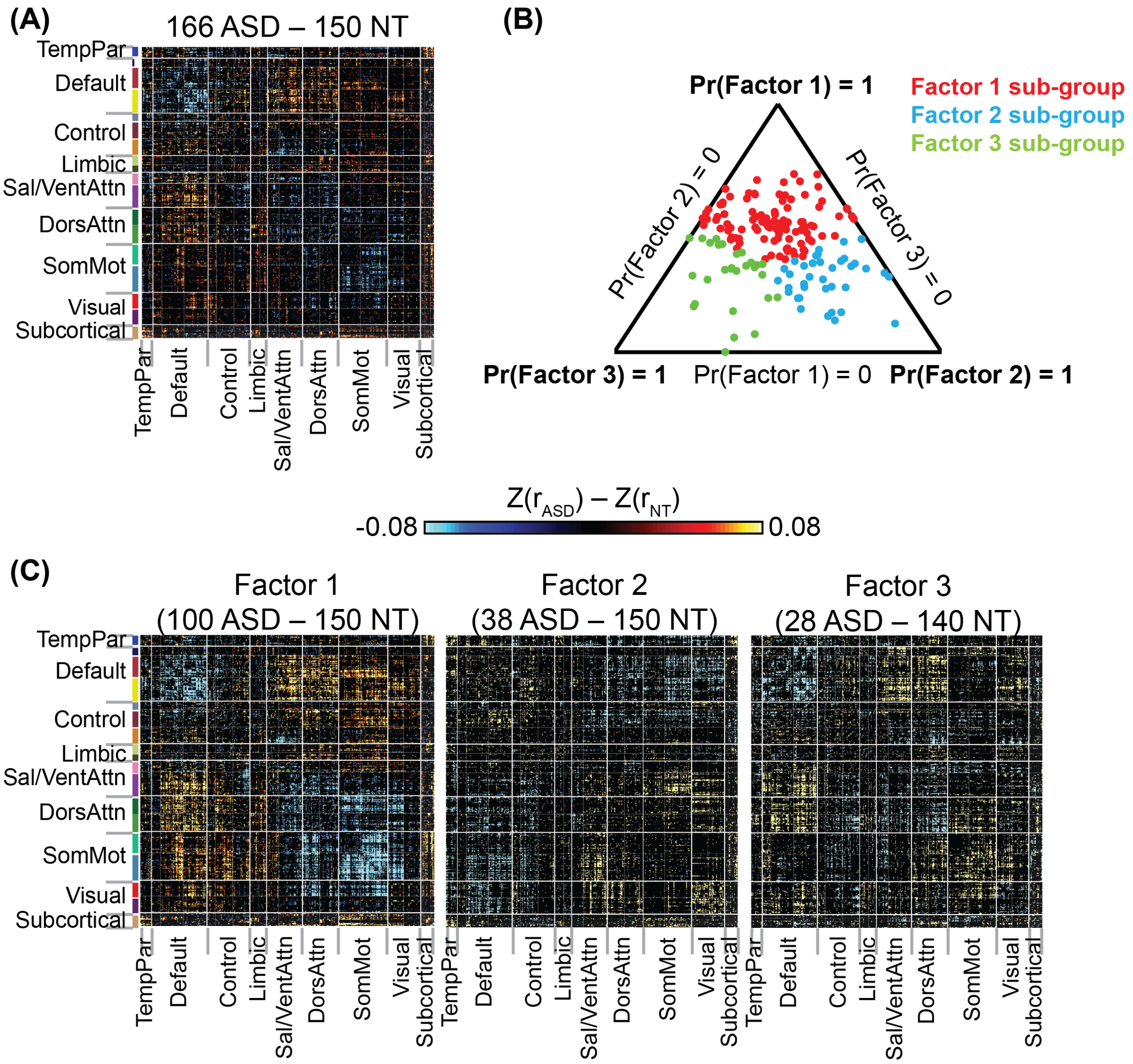


**Figure S6. Traditional case-control analyses (second subgrouping criterion) yielded weaker effects, while missing significant ASD-related RSFC associations.** (A) Traditional case-control analysis: RSFC differences between 166 ASD and 150 NT participants from ABIDE-I matched for age, sex and head motion within each site. ROI pairs with t-statistic > 2 are colored. RSFC differences were statistically significant after correcting for multiple comparisons (NBS $p < 0.0001$). Hot (cold) color indicates hyper-connectivity (hypo-connectivity) in ASD compared to NT participants. (B) Factor compositions of ABIDE-I ASD participants inferred using model parameters estimated from ABIDE-II+GENDAAR. Each participant corresponds to a dot, whose location (in barycentric coordinates) represents the factor composition. Corners of the triangle represent pure factors; closer distance to the respective corner indicates higher probability for the respective factor. Each ASD participant was assigned to the corresponding factor with highest probability Pr(Factor | Participant). Red, blue and green dots represent participants assigned to factor 1 (N = 100), factor 2 (N = 38) and factor 3 (N = 28) respectively. (C) RSFC differences between ASD and matched NT participants in the three groups (NBS $p < 0.0001$ for all three analyses). Effects were significantly stronger in sub-group analyses compared with case-control analyses despite the smaller sample sizes. Furthermore, case-control analysis showed similar hypo/hyper RSFC patterns as factor 1 ASD participants ($r$ = 0.68), but missed out on RSFC differences shown by participants assigned to factors 2 ($r$ = 0.23) and factor 3 ($r$ = 0.36).
